# Selective gating of neural modulation through frequency- and behavior-dependent modes during cortical electrical stimulation

**DOI:** 10.64898/2026.08.19.745734

**Authors:** Irene Rembado, Soo Yeun Lee, Lydia C. Marks, Leslie D. Claar, Areg Peltekian, Christof Koch, Costas A. Anastassiou

## Abstract

Electrical stimulation is widely used to modulate neuronal activity, yet its effects on neuronal circuits *in vivo* remain poorly understood. This, in turn, has hindered the principled design of stimulation protocols and raised questions about reproducibility that constrain the field’s translational impact. Here we combine cortical sinusoidal electrical stimulation (sES) with Neuropixels recordings to characterize stimulation-driven responses in more than 2,700 well-isolated neurons across 53 brain areas in 14 behaving, head-fixed mice. We uncover two distinct, concurrent modes of neural modulation. First is a sustained, brain-wide spike-phase entrainment effect that depends on stimulation frequency: entrainment to slow stimulation is supported by non-synaptic electric field propagation while anatomical connectivity dominates entrainment to higher stimulation frequencies. Second, we find a transient, spatially localized spike-rate modulation mainly mediated through anatomical connectivity that only emerges at high stimulation frequencies by selectively recruiting inhibitory neurons. We show that the two distinct modes are differentially shaped by behavior. By identifying how stimulation frequency governs the mechanism of neural engagement and how behavioral state selectively gates brain-wide entrainment but not local inhibitory recruitment, our results provide a mechanistic foundation for designing targeted, reproducible neuromodulation strategies.

## Introduction

Electrical stimulation has been used to modulate the nervous system for centuries, with the invention of electroshock by Cerletti and Bini in 1938^1^ marking the first modern therapeutic application of direct brain stimulation. Since then, electrical brain stimulation—from the invasive deep brain stimulation for Parkinson’s disease to the non-invasive transcranial alternating current stimulation for depression—has become a cornerstone of modern neurological and psychiatric therapy^2–5^. Among contemporary approaches, sinusoidal (continuous) electric stimulation (sES) has emerged as a particularly powerful and flexible tool. In humans, sES has been shown to modulate attention^6^, perception^7^, and memory^8^ and reduce symptoms in neurological and psychiatric disorders characterized by abnormal oscillatory activity^9^.

Early work established that the most excitable elements in cortical gray matter are myelinated axons of pyramidal neurons, with activation scaling with distance from the electrode tip ^10–13^. At the cellular level, thresholds and response polarity vary across cell types due to differences in morphology, laminar position, and connectivity ^14–16^. At the systems level, stimulation can modulate distant cortical regions via direct ortho- and anti-dromic axonal propagation as well as synaptic connections^17–21^. Even weak micro-stimulation activates sparse, spatially distributed neurons primarily by engaging axons of passage ^22^, with brief excitation often followed by prolonged suppression that shapes downstream propagation ^23–26^.

Mechanistically, sES differs from more conventional pulse stimulation protocols with its smoother temporal profile, modulating neuronal activity in a periodic manner while concurrently competing with ongoing oscillatory brain activity ^13,27–33^. The ability to interact with ongoing activity (as opposed to directly suppressing or exaggerating it) means that sES can effectively shape neuronal activity *in vivo* even at modest amplitudes. Yet, despite its promise and decades of clinical use showing clear therapeutic benefit, a fundamental challenge persists: identical stimulation parameters can produce different, or even opposite, neuronal effects across individuals and conditions ^32,33^. This variability has hindered the rational design of stimulation protocols and raised questions about reproducibility that constrain the field’s translational impact ^5,34–37^. In broader context, electrical stimulation effects remain difficult to interpret causally because they depend on the interaction among stimulation parameters, local cytoarchitecture, anatomical connectivity, and ongoing brain state ^14,22,38–42^. We therefore still lack a mechanistic framework that explains how these factors jointly determine circuit- and system-level responses across the brain.

Here, we provide the first brain-wide characterization of how electrical stimulation modulates neural activity *in vivo*. By combining cortical sES with large-scale Neuropixels 1.0 recordings in 2,725 well-isolated neurons across 53 brain areas in 14 behaving, head-fixed mice, we uncover two distinct, concurrent modes of neural modulation arising from separable biophysical mechanisms. The first is a sustained, ipsilateral brain-wide spike-phase entrainment whose underlying mechanism shifts with stimulation frequency — from ephaptic, field-driven coupling at low frequencies to connectivity-mediated, synaptic propagation at high frequencies. The second is a transient, spatially localized spike-rate modulation that emerges only at high stimulation frequencies through the selective recruitment of inhibitory neurons. These two modes differ not only in their spatial extent and temporal dynamics but also in their sensitivity to behavioral state: locomotion strongly modulates phase entrainment while leaving rate modulation unaffected. Together, these findings establish a mechanistic framework for understanding and predicting how electrical stimulation engages neural circuits across the brain.

## Results

### sES induces brain-wide spike-phase entrainment shaped by connectivity and geometry in a frequency-dependent manner

We applied bipolar sinusoidal electrical stimulation (sES) to the visual cortex (VIS) of 14 awake, head-fixed mice free to rest or run on a moving wheel (Fig. 1a). To characterize both proximal (<500 µm) and distal (<5 mm) effects of sES, we performed simultaneous ipsilateral Neuropixels recordings ^43^ inserted across several cortical and subcortical regions (Fig. 1b, S1). We systematically varied sES parameters (current intensity: 1 µA and 5 µA; frequency: 8 Hz, 28 Hz, and 140 Hz), with each sES protocol delivered 10 times (10 s stimulation epochs separated by 10-s inter-stimulation intervals). The oscillatory sES electric field robustly modulated the local field potential (LFP) across Neuropixels channels (Fig. 1c) resulting in a prominent spectral peak centered at the stimulation frequency (Fig. 1d). The spatial profile of the extracellular voltage amplitude reflected the bipolar stimulation through an asymmetric, double-peaked profile close to the electrode (Fig. 1e). The spatial amplitude profile of sES can be explained by the geometry of the experiment and relative positioning between the sES electrode and recording probes (Fig. 1f). That is, our spatial sampling covers many regions, with areas proximal to stimulation experiencing a bipolar stimulation profile whereas distant ones see a spatially diffuse far-field profile.

**Figure 1.**
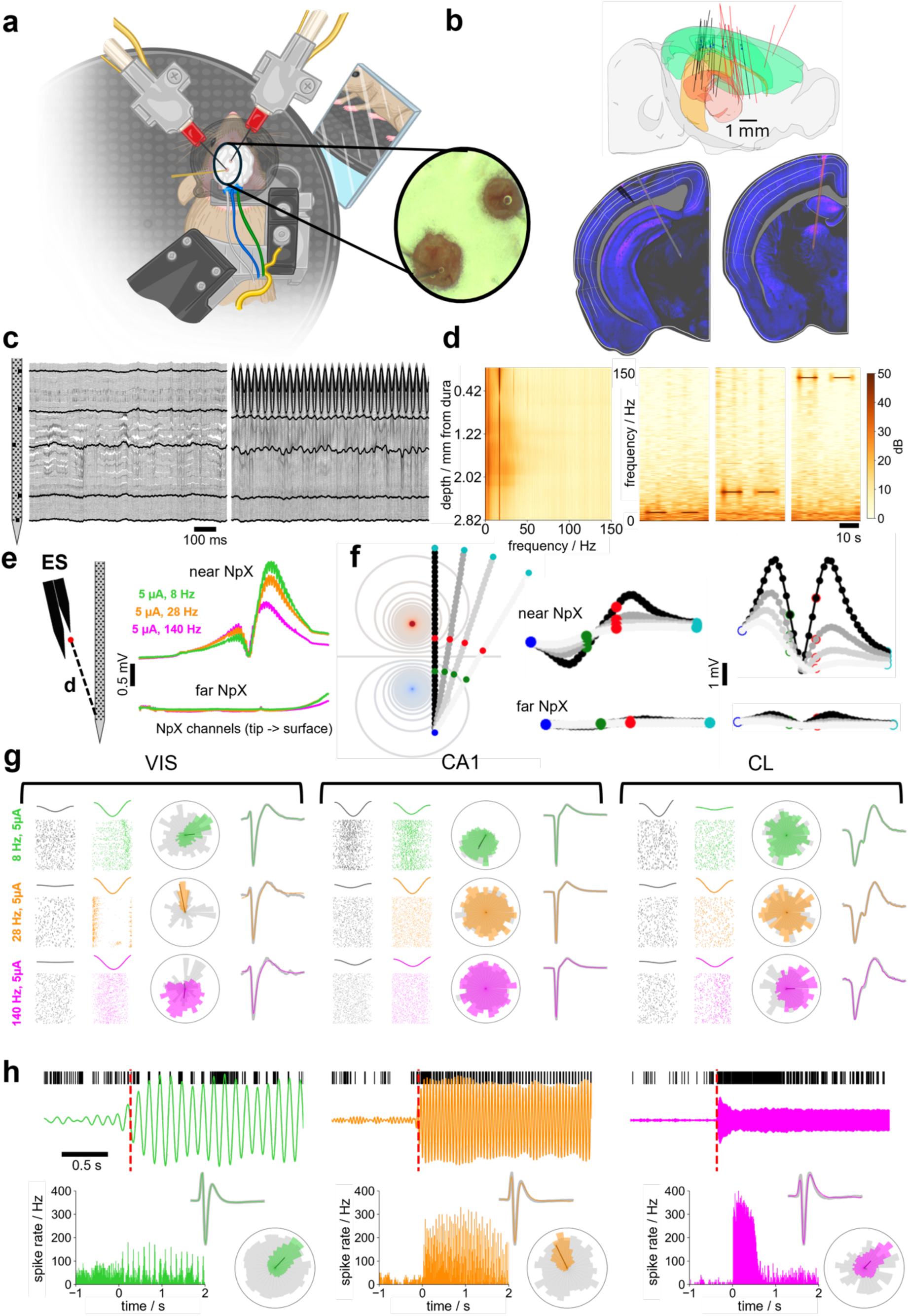
Brain-wide effects of sES delivered to the visual cortex of head-fixed mice. a) Schematic representation of the experiment. The mice are head-fixed and free to run or rest on a rotating wheel. Two Neuropixels 1.0 probes are inserted into two separate craniotomies, one exposing the visual cortical areas and the other targeting retrosplenial, somatosensory and motor cortical areas. A bipolar ES electrode is inserted into the visual craniotomy (inset: microscope image from one experiment). Pupil diameter is continuously measured from the right eye. b) Top: Neuropixels insertions across mice (*n*=14) shown using the reconstructed common coordinate framework (black: probe close to ES; red: probe far from ES; circle pairs: locations of bipolar ES electrodes). Below: Histological images of coronal brain slices showing the location of the bipolar ES electrode labeled in black (left, spanning visual area, layer 5) and the two Neuropixels probes (right, spanning layers of cortical areas, hippocampus, thalamic nuclei) with fluorescent dyes (pink). c) Representative 1 s LFP traces across Neuropixels channels during baseline (left) and during 28 Hz sES at 5µA (right; black: traces at specific locations; local field potential signal band: 0-1250 Hz). d) Left: power spectrum across all channels ordered as function of distance from pia in the absence of sES (probes inserted at an angle relative to pia). Right: time-frequency spectrum maps of one channel for sES at 8 Hz, 28 Hz, and 140 Hz sES, respectively, and 5µA. e) Left: schematic of the stimulating electrode position relative to the Neuropixels probe. Right: profile of the LFP-amplitude across the Neuropixels channels during sES (location relative to the sES electrode; top: probe near to sES electrode; bottom: far probe). f) Simple model of the LFP-amplitude for a bipolar ES in a purely resistive medium (point source approximation). Angle of the probe vs. stimulation electrode axis dictates LFP-amplitude symmetry (gray scale: probe angle). g) Spike-phase entrainment to sES for three units in primary visual cortex (VIS), hippocampus (CA1), and central lateral thalamus (CL). Entrainment effects for 8 Hz (green), 28 Hz (orange), 140 Hz (magenta) sES (amplitude: 5 µA). Left to right: raster plots at baseline (gray) and during sES (colored) showing the unit’s spike times relative to a single cycle of the LFP from a local electrode (see Methods); rose plot of the spike phase distribution for the unit across the whole recording at baseline (gray) and during sES (colored; black line: population vector during sES; overlap of spike unit waveforms during baseline (gray) and sES (colored). h) Spike rate effect to sES for a VIS unit for 8Hz (green), 28 Hz (orange) and 140 Hz (magenta) (sES amplitude: 5 µA). From top to bottom: single trial raster plot around sES onset ([-1, 2] s, broken red line: sES onset) overlapped to the LFP; inset: spike waveform overlap (control vs. sES); peristimulus time histograms (PSTHs) across ten sES repetitions (population vector as in panel g).

In its simplest form, the sES exerts a common, oscillatory signal to neurons, giving rise to a periodic membrane polarization that promotes spiking activity during the depolarizing phase and reduces it during the hyperpolarizing phase. One manifestation of such coupling between the periodic sES and neural activity is spike-phase entrainment where entrained neurons align to specific phases of the stimulation, while the overall spike rate remains broadly unaltered ^41,42^. When we looked at spike-phase entrainment, we saw that units drastically altered their spike phase distribution across brain areas and stimulation parameters (Fig. 1g). Another manifestation of electrical stimulation impacting neural activity is firing rate modulation, where stimulation onset and delivery cause a substantial change, either increase or decrease, of unit activity ^23,26,44^. We found many units showed transient firing rate modulation to sES during the initial period of stimulation delivery (Fig. 1h). sES therefore leads to two modes of neuronal activity coupling, spike-phase entrainment and firing rate modulation, which are concurrent but also vastly different in their spatiotemporal properties.

We first looked into spike-phase entrainment as a response to sES. To assess how sES coordinates spike timing, we calculated the instantaneous phase of LFPs using the Hilbert transform (see Methods). For each unit the spike-phase entrainment was then quantified using the population vector length (VL), where VL = 1 indicates perfect phase locking of a neuron to a particular sES phase and VL = 0 complete phase decorrelation. For each unit, we calculated VL during baseline (pre-stimulation) and during sES delivery, retaining only units that met spike waveform variability and excitability criteria under both conditions (waveform distortion score <0.1 compared to baseline and minimum of 51 spikes per condition, respectively; see Methods). Of the 53 brain areas sampled across all recordings, 27 met minimum inclusion criteria (≥8 units satisfying waveform stability and spike count thresholds; see Methods and Fig. S1) and were retained for subsequent analyses. We quantified spike–phase entrainment effects across these areas for different sES parameters and found that stimulation resulted in robust and sES parameter-specific spike-phase entrainment across the brain (Fig. 2a). Expectedly, stronger sES stimulation (5 µA) affected a larger fraction of units across areas compared to weaker sES (1 µA; Fig. 2a, compare two top rows). Moreover, units in areas closer to the sES site exhibited stronger spike-phase entrainment than farther away ones (Fig. 2a), though the distance scaling of the spike-phase entrainment showed sES frequency-specificity: lower frequencies (8 and 28 Hz) induced significant entrainment across a greater number of brain regions compared to high frequency sES (140 Hz). It follows that both stimulation frequency and current intensity influence the spatial extent of sES entrainment (Fig. 2a-b). Notably, these brain-wide effects occurred without accompanying changes in overall excitability, as firing rates remained stable throughout the 10-second stimulation epoch (Fig. S2). We conclude that sES gives rise to strong, brain-wide spike-phase entrainment while leaving firing rate broadly unaltered, with sES parameters dictating the spatial reach and overall constellation of this entrainment.

**Figure 2.**
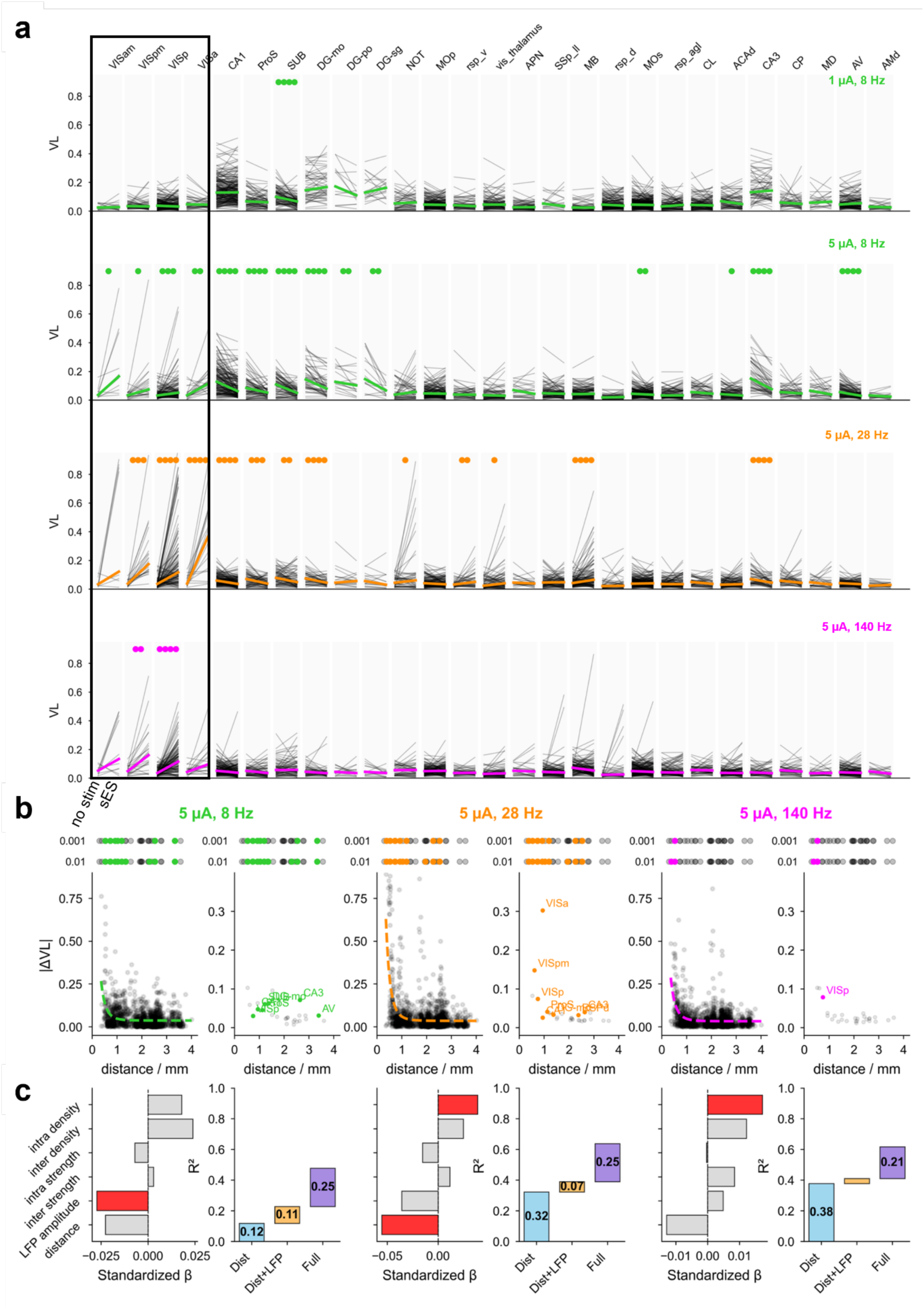
sES elicits brain-wide phase entrainment that propagates via anatomical connectivity and geometry in a frequency-dependent fashion across distributed brain networks. a) Spike-phase entrainment measured by population vector length (VL) compared between baseline (no stim) and sES across 27 brain areas for four sES conditions (top to bottom; lines connect the same unit between baseline - left - and sES - right). Statistical significance of VL difference above each area (circle color corresponds to sES frequency; number of circles indicates significance level: ●●●● p ≤ 0.0001; ●●● p ≤ 0.001; ●● p ≤ 0.01; ● p ≤ 0.05; paired t-test on pairwise differences FDR-corrected). Only units satisfying minimum activity criteria during baseline and sES included (row 1: n = 2,275; rows 2–4: n = 1,958 units). Brain areas ordered by increasing distance from sES location (left to right), with unit position estimated from the Neuropixels electrode. Black box: visual cortical areas adjacent to sES electrode. b) Spatial decay of phase entrainment as a function of distance from the sES site for 8, 28, and 140 Hz (5 µA; n = 1,958 units across 27 areas per frequency). Circles above each panel indicate area-level significance of entrainment (colored: significant; black: non-significant) at two thresholds (upper row: p < 0.001; lower row: p < 0.01; one-sample t-test on within-area VL differences). Left panels: |ΔVL| (|VL_stim − VL_pre|) for individual units plotted against distance from the sES site. Solid colored line: fitted inverse power law with offsets, f(d) = c + 1/(d − d_0_)ⁿ, where d is distance in mm. Right panels: area-averaged |ΔVL| plotted against mean area distance; colored circles with area labels denote significantly entrained areas (p < 0.001); black circles denote non-significant areas. Spatial decay is frequency-dependent, with lower frequencies (8 Hz) showing shallower decay and longer-range effects than higher frequencies (140 Hz). c) Hierarchical regression analysis quantifying contributions to entrainment strength across brain areas (areas excluded if missing connectivity data). Left panels: standardized regression coefficients (β) from full linear model including distance from sES, LFP amplitude at sES frequency, and four anatomical connectivity metrics derived from the Allen Mouse Brain Connectivity Atlas ^45,46^ (inter-area projection strength, intra-area connection strength, inter-area projection density, intra-area connection density). All predictors z-scored before regression. Red bars: significant predictors (p < 0.05, two-sided t-test); gray bars: non-significant predictors. Right panels: cumulative R^2^ from hierarchical regression. Model 1 (blue): distance alone. Model 2 (orange): distance + LFP amplitude. Model 3 (purple): full model including connectivity metrics. Numbers indicate ΔR^2^ contributed by each model step. The β coefficients and the hierarchical ΔR^2^ provide complementary mechanistic information. The β’s identify which *individual* predictors carry the most weight after controlling for all others, while the ΔR^2^ quantifies the *collective* variance explained by each predictor block. The identity of the strongest individual predictor shifts with frequency: at 8 Hz, LFP amplitude dominates (consistent with ephaptic, field-based coupling), whereas at 140 Hz, intra-area connectivity density is the leading predictor (consistent with synaptic, circuit-mediated coupling). Despite this shift in the dominant individual predictor, connectivity metrics collectively explain 21–25% additional variance beyond geometric factors across all frequencies (ΔR^2^ = 0.25, 0.25, and 0.21 for 8, 28, and 140 Hz, respectively), demonstrating a consistent and substantial contribution of anatomical network architecture to entrainment propagation at every frequency tested.

We next asked how circuit properties impact spike phase-entrainment and shape the overall circuit’s response to the stimulation. To identify the factors governing the spatial pattern of spike-phase entrainment, we performed hierarchical linear regression on area-level entrainment strength (|ΔVL|) using three nested models of increasing complexity (Fig. 2c). The first model included distance from the stimulation electrode alone; the second added local field potential (LFP) amplitude at the stimulation frequency; the third incorporated four anatomical connectivity metrics derived from the Allen Mouse Brain Connectivity Atlas ^45,46^: inter-area projection strength and density, and intra-area connection strength and density. The regression revealed two key findings.

First, the identity of the strongest individual predictor shifted systematically with stimulation frequency. At 8 Hz, LFP amplitude was the dominant single predictor of entrainment strength (standardized β; Fig. 2c, left panels) consistent with ephaptic coupling, in which the amplitude of the exogenous electric field at each brain area determines the strength of non-synaptic modulation. At 140 Hz, this pattern reversed: intra-area connectivity density emerged as the strongest individual predictor, consistent with synaptic-mediated coupling in which local network architecture governs the stimulation response. At 28 Hz, the pattern was intermediate, with distance as the leading predictor and both field- and connectivity-based factors contributing.

Second, despite the shift in the dominant individual predictor (as measured by the standardized β), anatomical connectivity metrics collectively explained 21–25% of additional variance beyond what geometric factors (distance and LFP amplitude) could account for (ΔR^2^ = 0.25, 0.25, and 0.21 for 8, 28, and 140 Hz, respectively; Fig. 2c, right panels). This collective contribution remained stable across frequencies, even as the relative importance of individual connectivity metrics changed. At 8 Hz, where distance alone explained only 12% of the variance, the addition of connectivity more than doubled the full model’s explanatory power (total R^2^ = 0.48). At higher frequencies, distance contributed more substantially (R^2^ = 0.32 and 0.38 for 28 and 140 Hz), but connectivity still added a large and consistent increment. To control for coupling due to more complex spike-phase entrainment profiles (e.g. multimodal spike-phase distributions as a result of field entrainment), we also looked at an alternative metric, the mean modulation ratio (Fig. S3), that accounts for such effects and found very similar results (Fig. S4). Note that the discrepancy between the individually modest β coefficients of single connectivity metrics and their large collective ΔR^2^ reflects the intercorrelation among the four connectivity measures: each capture partially overlapping aspects of anatomical network organization, suppressing individual standardized coefficients while preserving their joint explanatory power.

We wondered whether the observed entrainment effects are the result of sES in visual cortex only or whether similar effects are observed when stimulating adjacent cortical regions. For a subset of the animals, post-hoc histological analysis revealed the stimulating electrode tip to be in somatosensory cortex (n=4) or secondary motor cortex (n=1). For these cases we saw the same brain-wide spike-phase entrainment as when administering sES into visual cortex. Taken together, these results demonstrate a frequency-dependent mechanistic transition: low frequency entrainment is best predicted by the LFP amplitude (consistent with ephaptic propagation), whereas high-frequency entrainment is best predicted by local circuit connectivity (consistent with synaptic propagation). At all frequencies, anatomical connectivity contributes substantially to entrainment beyond what spatial geometry alone can explain, establishing that network architecture is a fundamental determinant of how sES engages across brain areas.

Given brain-wide spike-phase entrainment, we next asked whether all neurons in an area homogeneously respond to sES by collectively increasing their spike-phase coupling or whether there exist communities within each area that preferentially phase-lock. To determine whether sES recruits distinct neuronal subpopulations, we analyzed single-unit spike–phase entrainment at baseline (no stimulation) compared to during sES (Gaussian mixture modeling, optimal number of clusters selected by maximizing the silhouette score). At 28 Hz and 5 µA, a stimulation regime where both geometry and connectivity affect entrainment (Fig. 2c), units segregated into distinct clusters across highly entrained brain areas (Fig. 3a). Importantly, such clustering was observed not only in areas near the stimulation site but also at more distal locations with heterogeneous entrainment responses emerging across the brain areas we recorded from (Fig. 3b). Moreover, heterogeneous within-area spike-phase entrainment was robust and occurred across stimulation frequencies (Fig. 3c). Analysis across stimulation frequencies revealed significant clustering in most recorded regions, with 94% of the 27 retained brain areas (constraints to include an area for GMM analysis: units per area > 8) exhibiting significant clustering at one or more frequencies (FDR-corrected *p* < 0.05; Fig. 3c). It follows that while spike-phase entrainment occurs across the entire brain, there exists significant heterogeneity within the units of each area in their coupling to the imposed sES.

**Figure 3.**
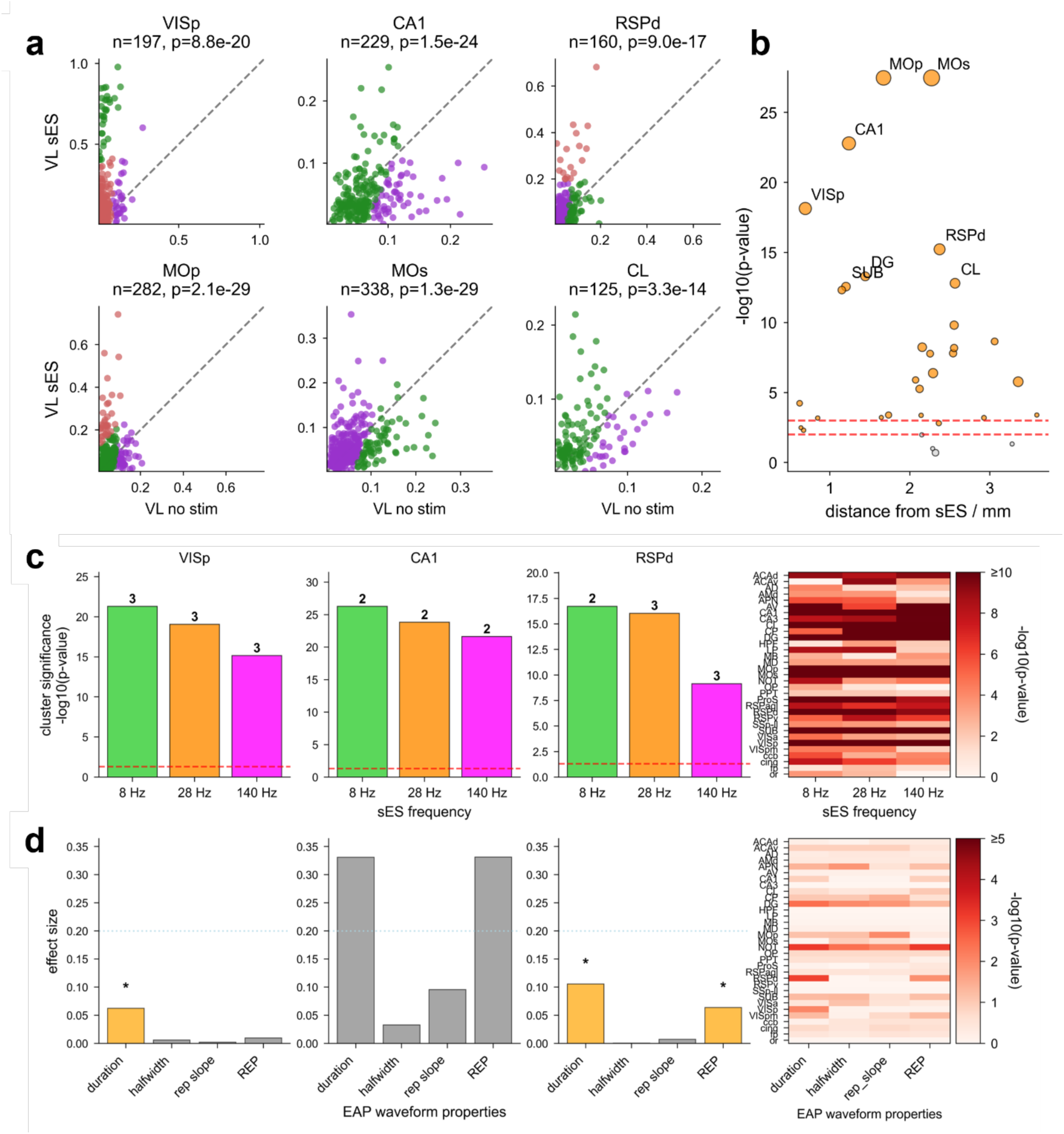
sES recruits distinct neuronal subpopulations independent of cell-type identity. a) Comparison of spike-phase entrainment during baseline (no stim, on the x-axis) vs. sES for individual units (on the y-axis) in six representative brain areas at 28 Hz and 5 µA. Colors: cluster membership determined by Gaussian mixture modeling. Circles above the diagonal represent neurons with enhanced phase entrainment. All areas show highly significant VL increases during sES (Wilcoxon signed-rank test; *n* and *p*-values indicated). b) Clustering significance (−log_10_(FDR-corrected p-value)) plotted against distance from the sES site across all recorded areas. Colored circles indicate areas with significant clustering (FDR-corrected p < 0.01); gray circles indicate non-significant areas. Circle size is proportional to the number of recorded units per area. Dashed red horizontal lines mark FDR-corrected p = 0.01 and p = 0.001 thresholds. Strong clustering occurs at both proximal and distal sites, consistent with connectivity-driven rather than distance-dependent propagation of entrainment. c) Left: Clustering significance across sES frequencies (8, 28, 140 Hz) for VISp, CA1, and RSPd (numbers: optimal cluster count). Right: Heatmap summarizing clustering significance (−log_10_(p-value)) for all recorded areas. Of 27 areas analyzed, 94% showed significant clustering at one or more frequencies (FDR-corrected p < 0.05), demonstrating that heterogeneous entrainment responses are a general feature of brain-wide stimulation. d) Left: Effect sizes (η^2^) for associations between four EAP waveform properties (duration, half-width, repolarization slope, REP) and cluster membership in VISp, CA1, and RSPd (left to right) at 28 Hz. Dashed line indicates small effect size threshold (0.2). Orange bars denote FDR-corrected p < 0.05 (Mann-Whitney U test for 2-cluster areas; Kruskal-Wallis test for ≥3 clusters; corrected across features); gray bars: non-significant associations. Asterisks: FDR-corrected p < 0.05. Right: heatmap of −log_10_(FDR-corrected p-value) for each waveform property across all recorded areas. Effect sizes are uniformly small (all below 0.2) even for significant associations, indicating that waveform properties have negligible predictive power for cluster identity. Dissociation points to clusters that reflect frequency-specific functional network states rather than stable cell-type differences.

Where does this heterogeneity in the within-area spike-phase entrainment response to sES come from? One explanation could be that since each area comprises of diverse cell types, heterogeneity in entrainment reflects different classes or types responding to the electrical stimulus in a distinct manner. The most prominent such distinction is between excitatory and inhibitory neurons with excitatory neurons having a typically elongated dendritic morphology compared to the more symmetric inhibitory morphologies with the former associated with more pronounced field effects compared to the latter ^14,16,47^. Excitatory and inhibitory cortical units typically separate based on their extracellular spike waveform features with excitatory units being associated with wider (longer duration, halfwidth, etc.) waveforms whereas inhibitory units with more narrow ones. We tested whether associations between cluster membership and extracellular action potential waveform properties explain the entrainment clustering and found that waveform features only weakly related to cluster identity with only 6.1% of area–property combinations exceeding both statistical significance and moderate effect size thresholds (η^2^ > 0.2; Fig. 3d). It follows that spike-phase entrainment clusters do not reflect stable cell-type classifications (e.g., regular- vs. fast-spiking units), but instead likely correspond to frequency-specific functional networks dynamically engaged and configured by sES.

### sES induces a transient, local, sES-frequency and cell-type specific spike-rate modulation

We next looked at how sES modulates neural firing rate. On the one hand, units did not show a change in excitability over the 10 s duration of sES delivery (Fig. S2). However, a subset of units responded with strong and transient firing rate modulation at the onset of sES (Fig. 1c, 4a) which became more pronounced and lasted longer with increasing stimulus frequency, peaking at a median duration of 270 ms at 140 Hz (Fig. 4a-b). Specifically, we identified two distinct onset responses: a transient firing rate increase (> 1.1 × baseline) and a transient rate decrease (< 0.95 × baseline). In stark contrast to spike-phase entrainment, the transient firing rate modulation was not brain-wide but, instead, was spatially confined close to the sES delivery location (Fig. 4b). These effects were only observed for 5 µA, with 1 µA not producing a measurable spike-rate modulation (Fig. S5). Moreover, the observed firing rate changes were independent from the brain-wide spike-phase entrainment (Fig. S6). We conclude that parallel to the brain-wide spike-phase entrainment, sES also produces a highly localized (close to the stimulation electrode), transient (less than 300 ms from stimulus onset), and stimulus frequency-specific firing rate-modulation.

**Figure 4.**
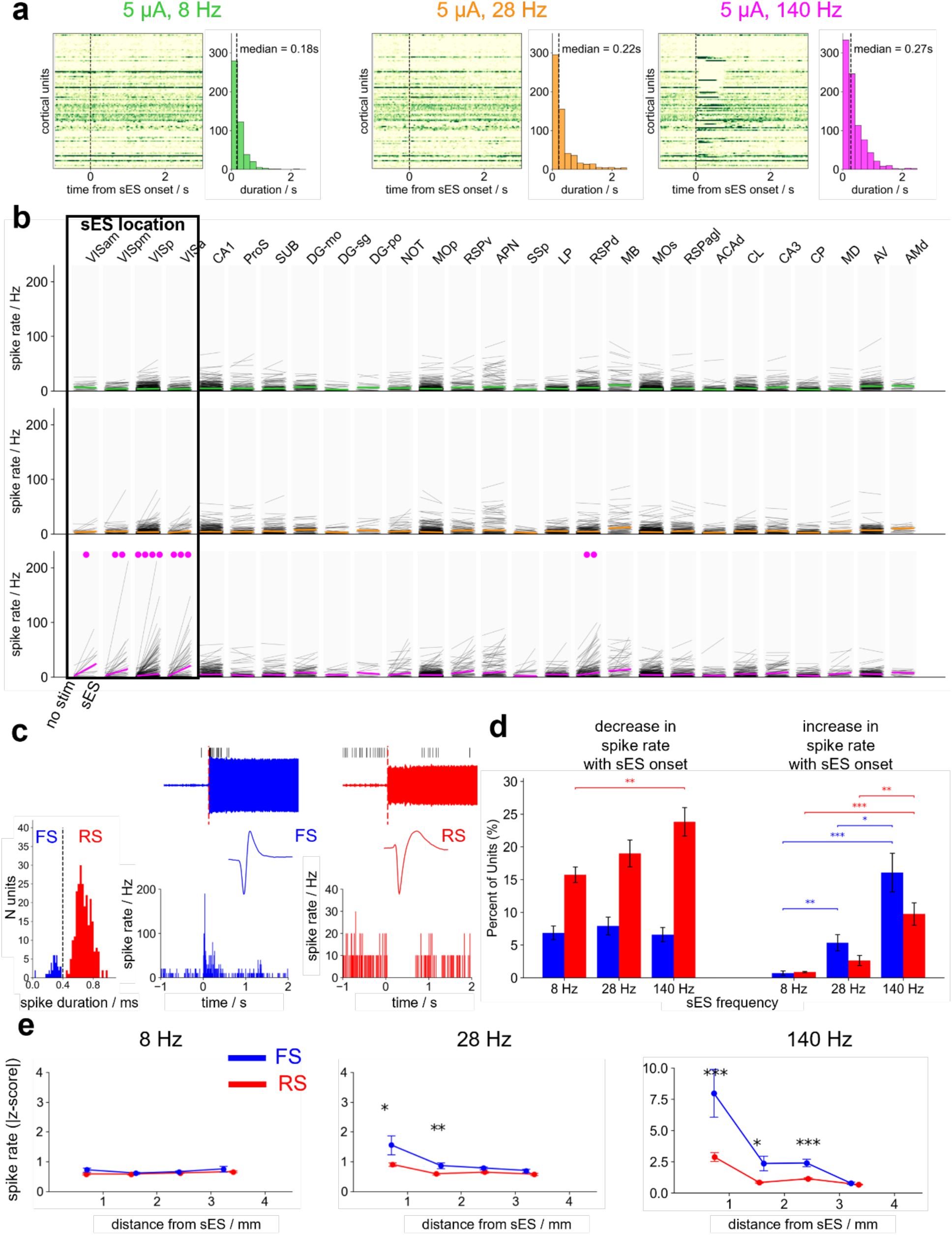
Local bipolar stimulation induced frequency-specificity, cell-type dependent and localized transient firing rate changes. a) Spike density function of the evoked firing rate across all trials for cortical units near E-Stim for one representative mouse as function of time from stimulation onset across protocols (green color map left to right: 5 µA at 8 Hz, 28 Hz, 140 Hz, respectively) with distribution of the duration of the transient effects across protocols at 5 µA. Vertical black line shows the median value for each distribution. b) Average firing rate for 100 ms time window after stimulation onset compared to baseline pre-stimulation across brain areas and protocols (top to bottom: 5 µA at 8, 28, and 140 Hz, respectively). Significance assessed via paired t-test on within-unit firing rate differences with Benjamini-Hochberg FDR correction (*p < 0.05, **p < 0.01, ***p < 0.001, ****p < 0.0001). No such effects were found at 1 µA (Fig. S5). c) Representative example of a regular-spiking unit (RS, red) and a fast-spiking unit (FS, blue) responses at the onset of 140Hz sES at 5 µA. Left: Distribution of the spike waveform duration with the threshold of 0.4 ms discriminating RS from FS. Right: For each unit from top to bottom: single trial raster plot around three seconds from stimulation onset [-1, 2] s overlapped to its peak channel LFP signal; spike waveform; peristimulus time histograms (PSTHs) across all ten stimulation sweeps. d) Percent of RS (red bars) and FS (blue bars) that show a transient decrease or increase of firing rate across protocols at 5 µA calculated across animals (mean ± SE; pairwise Mann Whitney test was computed for each cell type across stimulation protocols for each type of transient effects, \**p* < 0.05, \*\**p* < 0.01, ***p< 0.001 and **** p<0.0001). e) Absolute Z-scored firing rate relative to baseline pre stimulation during the first 100 ms of stimulation for all RS and FS units as function of their distance from the stimulating electrode (E-Stim) across protocols (top to bottom: 5 µA at 8 Hz, 28 Hz, 140 Hz, respectively). Statistical testing: Mann-Whitney with Benjamini-Hochberg multiple comparison correction. Statistical significance: * p<0.05; ** p<0.01, *** p<0.001 and **** p<0.0001.

Given the characteristics of the firing rate modulation, we wondered whether it is supported by distinct cell classes. In this case, we separated into putatively excitatory regular-spiking (RS) and putatively inhibitory fast-spiking (FS) (Fig. 4c) and quantified the portion of RS and FS increasing or decreasing their firing rate for each animal (Fig. 4d). We found that increasing stimulation frequency resulted in the preferential and increasing fraction of FS transiently increasing their firing rate (FS fraction increasing spike rate at 8 Hz: <1%; at 140 Hz: 16%; Fig. 4d). In contrast, increasing stimulation frequency exerted a more modest effect on RS (RS fraction increasing spike rate at 8 Hz: <1%; at 140 Hz: 9.7%; Fig. 4d). What about the units transiently reducing their spike rate? We found that increasing stimulation frequency resulted in a higher fraction of RS with significant firing rate decrease while for FS that fraction remained unchanged (RS with a spike rate decrease at 8 Hz: 15.7%; at 140 Hz: 23.8%; FS with a spike rate decrease at 8 Hz: 6.8%; at 140 Hz: 6.5%; Fig. 4d). These patterns support the frequency-dependent recruitment of local inhibition that, in turn, silences excitatory neurons with the fraction and balance between excitatory and inhibitory dictated by the sES frequency. Finally, we also asked whether distance from sES location differentially affected FS vs. RS. We found that sES preferentially affects FS over RS but that this effect is highly distance-specific with the firing rate effect on FS being more localized compared to RS at 140 Hz (at 28 Hz: *p*-value (regression slope comparison FS vs. RS) = 8.5 10^-4^; at 140 Hz: p-value (slope comparison FS vs. RS) = 8.4 10^-7^; Fig. 4e). In summary, the spatially localized effect exerted by sES is mainly attributed to the recruitment of local inhibitory neurons – their transient (200-300 ms) increase in firing rate strongly suppresses local excitatory neurons for fast but not slow stimulation frequencies.

### Behavioral state differentially affects spike-phase entrainment and spike-rate modulation induced by sES

Our data demonstrates that cortical sES acts like a common signal, synchronizing neuronal dynamics brain-wide through spike-phase entrainment as well as locally via transient firing rate modulation. At the same time, behavior can exert a powerful modulatory signal on visual cortex (e.g. ^48–51^). We therefore asked whether behavior interacts with spike-phase entrainment and firing rate modulation and whether one mechanism is more affected by behavior than the other. We looked at run vs. rest dynamics of head-fixed animals running freely on a wheel in the presence and absence of sES. For the spike-phase entrainment (spikes labeled as *rest* or *run* based on wheel speed) we found that the behavior significantly influenced entrainment both during baseline (no stimulation) and throughout sES epochs (Fig. 5a-b; Fig. S7). Importantly, the behavioral modulation of spike phase-entrainment also depended on sES parameters: behavior significantly affected spike-phase entrainment in a large fraction of brain areas for low sES frequencies, with that fraction decreasing for higher stimulation frequencies (77% at 8 Hz; 30% at 28 Hz; <10% at 140 Hz; Fig. 5c). In contrast, the fraction of brain areas exhibiting a change in firing rate-modulation between run and rest was substantially smaller (15% at 8 Hz; 15% at 28 Hz; 0% at 140 Hz; Fig. 5d-f). Notably neither stimulation onset nor offset elicited changes in locomotion or pupillometry (Fig. S8) indicating that the animals did not overtly respond to sES. We conclude that behavior affects the two coupling mechanisms distinctly: while brain-wide spike-phase entrainment is significantly impacted by running particularly for low stimulation frequency and remains broadly unaltered for high frequency, the transient firing rate modulation local to stimulation remains broadly unaffected by behavior.

**Figure 5.**
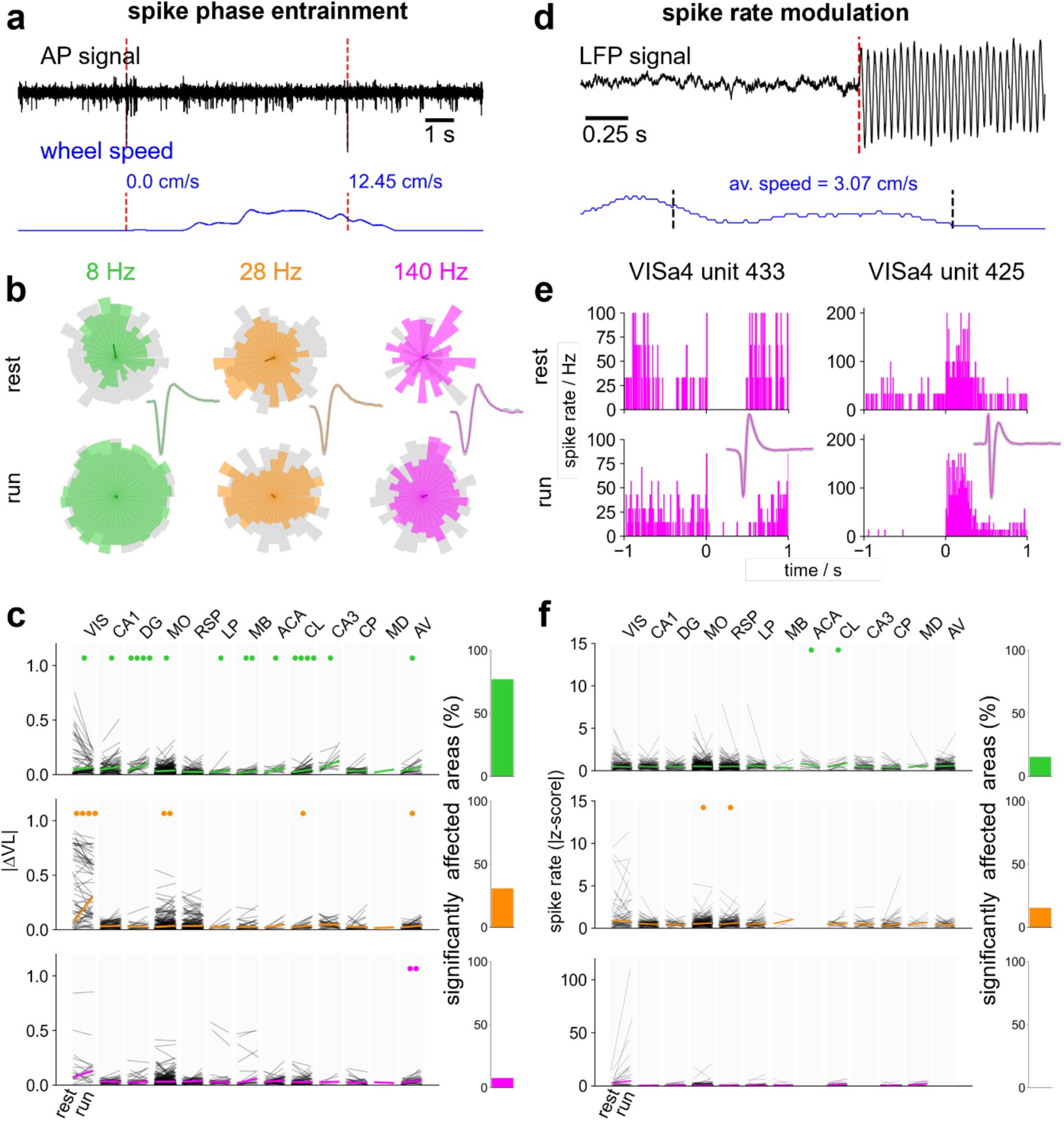
The behavioral state critically modulates spike-phase entrainment in a frequency- and area-dependent manner without impacting the spike-rate modulation. a) Spike-phase entrainment. Representative classification of a unit’s spikes (action potential raw signal shown in black) as “rest” or “run” depending on the value of the average wheel speed (blue) in a time window of 200 msec around the time of spike’s occurrence (red vertical lines). In this example the first spike occurred while the wheel speed was at 0 cm/s (i.e. the animal was resting), while the second spike occurred when the wheel speed was at 12.45 cm/s (i.e. the animal was moving). b) Rose plots of the phase distributions of a representative unit during resting (top) and during running (bottom) for baseline (gray) and during the sES across three stimulation protocols (8Hz, green; 28 Hz, orange; 140 Hz, pink at 5 µA). Spike waveform at baseline (gray) and during sES (color) is also shown. c) Length of the absolute value of population vector difference (|ΔVL|) between sES and baseline for rest and run across brain areas and protocols (top to bottom: 5 µA at 8 Hz, 28 Hz, 140 Hz, respectively). When unit satisfies the minimum requirements (*n* spikes> 51 for each condition and circular fit error < 0.1; see Methods) in both conditions, line is shown. Thick colored lines: median |ΔVL| within area for each condition. The behavioral states significantly affect the spike-phase entrainment brain-wide specifically at lower but not at higher frequencies. The bar plot on the right shows the percentage of significantly affected areas per sES protocol. d) Spike-rate modulation. Representative classification of stimulation onset as “rest” or “run” depending on the value of the average wheel speed (blue) in a time window of 1.5 sec around the stimulation onset (red vertical lines) (see Methods for details). e) Peristimulus time histograms (PSTHs) for two VISa4 units across all classified rest (top) and run (bottom) trials at the onset of 140 Hz sES at 5µA. Spike waveform at baseline (gray) and during sES (pink) is also shown. f) Absolute value of the z-scored firing rate calculated over 100 ms after stimulation onset relative to 1 s baseline pre stimulation for rest and running trials across brain areas and protocols (top to bottom: 5 µA at 8 Hz, 28 Hz, 140 Hz, respectively). When unit satisfies the minimum requirements in both conditions, line is shown. Thick colored lines: median z-scored firing rate within area for each condition. The bar plot on the right shows the percentage of significantly affected areas per sES protocol. Statistical testing: Mann-Whitney with Benjamini-Hochberg multiple comparison correction. Statistical significance: * p<0.05; ** p<0.01, *** p<0.001 and **** p<0.0001.

## Discussion

We provide a comprehensive brain-wide, *in vivo* characterization of how sinusoidal electric stimulation (sES) shapes neural activity across distributed ipsilateral circuits of the mouse brain. By combining cortical sES with brain-wide Neuropixels recordings, we uncovered two distinct, concurrent modes of neural modulation, spike-phase entrainment and firing-rate modulation, that arise from separable biophysical mechanisms, recruit different cell-clusters, occupy different spatiotemporal scales, and are differentially governed by behavioral state. These findings bridge controlled *in vitro* observations with circuit-level dynamics operating in the intact brain, showing that stimulation simultaneously engages at least two mechanisms whose relative contributions shift with stimulation parameters, local and brain-wide circuit properties, cellular excitability, and brain state.

Anatomical connectivity is a fundamental determinant of how spike-phase entrainment propagates across the brain. Connectivity metrics derived from the Allen Mouse Brain Connectivity Atlas ^45,46,52^ collectively explained 21–25% of the variance in entrainment strength beyond what geometric factors could account for, a contribution that remained consistent across all stimulation frequencies tested. Hence, beyond passive current spread, network propagation through anatomical wiring shapes the spatial pattern of entrainment ^10,12,22^, consistent with studies showing that stimulation-evoked activity often exceeds passive field spread^17^ and stimulation-evoked potentials follow anatomical pathways ^26,53–58^.

While the collective contribution of connectivity was consistent across frequencies, the regression analysis revealed that the mechanism underlying spike-phase entrainment shifts systematically with stimulation frequency. At 8 Hz, LFP amplitude, a measure of the exogenous electric field experienced by each brain area, was the strongest individual predictor of entrainment strength. This pattern is consistent with ephaptic coupling, whereby even weak but slow sES already potently entrains the neuronal membrane proportionally to the field amplitude ^27,41,59^. At 140 Hz, this pattern reversed: intra-area connectivity emerged as the dominant individual predictor, supporting that the local synaptic circuit architecture, not the amplitude of the exogenous field, governs entrainment at high frequencies. This frequency-dependent shift in the dominant predictor, from field amplitude (ephaptic) to circuit connectivity (synaptic), provides direct *in vivo* evidence for a mechanistic regime shift. Previous *in vitro* work established that endogenous ephaptic coupling is strongest at low frequencies for excitatory neurons ^27,41^ whereas inhibitory neurons tend to entrain to faster frequencies^42^. Here we show *in vivo* that ephaptic and synaptic mechanisms are not mutually exclusive modes but rather coexist along a stimulation frequency continuum, with their relative contributions shifting systematically with stimulation frequency. This mechanistic transition provides a concrete biophysical explanation for the well-documented frequency dependence of stimulation outcomes in both animal models and human clinical studies ^5^.

Importantly, within individual brain areas, neurons did not respond uniformly to sES but segregated instead into distinct response clusters characterized by different spike-phase entrainment profiles. This heterogeneity was pervasive, with 94% of brain areas showing statistically significant clustering across stimulation frequencies. This cluster consistency suggests that within-area response diversity is not random but reflects robust organizational principles. Surprisingly, these response clusters did not correspond to classical cell-type distinctions based on extracellular waveform properties: only 6% of area–waveform property combinations reached statistical significance. This, in turn, suggests that these clusters do not represent excitatory vs. inhibitory responses, at least not in a traditional sense. Instead, the data supports recent evidence ^13,60^ that entrainment clusters reflect neurons within functional networks like synapsembles ^61^ the composition of which is mainly determined by its local and long-range connectivity and intrinsic excitability rather than a specific cell class or type.

The second mode of modulation, a transient (<300 ms), spatially confined firing rate change at stimulation onset, stands in stark contrast to the sustained, brain-wide nature of spike-phase entrainment. This mode was largely absent at low stimulation frequencies and emerged with increasing frequency through the selective recruitment of fast-spiking inhibitory neurons, rising from less than 1% at 8 Hz to 16% at 140 Hz. The recruited inhibitory neurons, in turn, suppressed firing of nearby excitatory cells, consistent with prior observations that single-pulse electrical stimulation induce a brief excitation frequently followed by a prolonged inhibitory period that shapes downstream propagation ^23–26,44^. Yet, our findings extend these observations in two important ways. First, we show that this phenomenon is gated by frequency: it is absent at low frequencies and emerges progressively as the stimulation frequency increases. Second, the spatial restriction of firing rate modulation to areas near the stimulation electrode, coupled with its dependence on anatomical connectivity rather than distance alone, suggests that this mode reflects synaptic activation of local inhibitory circuits rather than a direct, non-synaptic effect on neuronal membranes. The selective engagement of fast-spiking neurons at high frequencies is reminiscent of the frequency ranges used therapeutically in deep brain stimulation for Parkinson’s disease, where high-frequency stimulation (140 Hz or higher) suppresses pathological activity through local inhibitory mechanisms ^37,62^. While our preparation differs substantially from clinical DBS, the convergence of frequency selectivity and inhibitory recruitment suggests that shared circuit-level principles may underlie both single-pulse electrical stimulation and high-frequency sES.

Another important observation is the dissociation between the spike-phase entrainment and firing-rate modulation with respect to behavioral state. Locomotion strongly modulated phase entrainment: 77% of brain areas showed significant behavior-dependent entrainment changes at 8 Hz, but this effect essentially vanished at 140 Hz (less than 10% of areas). In contrast, the transient rate modulation remained invariant across behavioral states at all frequencies tested. This dissociation has not been reported previously and has direct implications for the notorious variability of stimulation outcomes (e.g. ^34,36^). The brain state dependence of low-frequency phase entrainment is consistent with the known sensitivity of endogenous theta and alpha oscillations to arousal, attention, and locomotion ^50,63^. Because low-frequency entrainment acts predominantly through ephaptic coupling, shaped by the electrostatics of the brain tissue as well as the ongoing state of the network, it is expected to interact strongly with endogenous oscillations like hippocampal theta and neocortical beta. Conversely, the behavioral invariance of rate modulation, which operates through synaptic activation of local inhibitory circuits, may reflect the stereotyped input–output properties of fast-spiking interneurons, whose high firing rates, narrow temporal integration windows, and rapid, reliable firing make their recruitment and inhibitory output comparatively insensitive to slow fluctuations in network state ^64–66^. The implication is significant: stimulation protocols that rely on phase entrainment may produce state-dependent and therefore variable outcomes, while those engaging local inhibitory circuits at high frequencies may be more reproducible across behavioral conditions. This framework therefore addresses the variability problem that has hindered the rational design of stimulation protocols and raised questions about reproducibility ^36,38^. Furthermore, in a more clinical setting, it also provides a plausible explanation for the contradicting effects induced by 10 Hz stimulation shown to increase alpha power in schizophrenia but decrease it in major depressive disorder, conditions characterized by opposite baseline alpha abnormalities ^32,33^.

Taken together, our findings offer principled constraints for the design of stimulation protocols according to the intended therapeutic target. Low-frequency sES produces brain-wide, ephaptic-dependent phase entrainment that synchronizes distributed networks but is sensitive to behavioral state, making it potentially suitable for conditions characterized by pathological desynchronization across brain networks, such as the oscillatory abnormalities observed in schizophrenia and depression ^32,33,67^, provided that state-related variability can be managed through closed-loop or state-adaptive paradigms ^68–70^. High-frequency sES engages local inhibitory circuits in a behaviorally invariant manner, suggesting utility for conditions requiring spatially targeted, reproducible suppression of pathological activity, as in the high-frequency stimulation protocols used for movement disorders ^3,37^. Collectively, these results position sES as a circuit phenomenon governed by frequency-dependent shifts between field-driven and connectivity-mediated mechanisms, providing a mechanistic foundation for interpreting variability across brain states and experimental conditions. By revealing how frequency, connectivity, cellular excitability, and behavioral state jointly determine whether stimulation synchronizes distributed networks or recruits local inhibitory circuits, this work provides a principled roadmap for designing more precise and reproducible stimulation strategies.

## Author contributions

Conceptualization: C.A.A.; Formal analysis: I.R., S-Y.L., A.P., C.A.A.; Investigation: I.R., L.C.M, C.A.A.; Funding acquisition: C.A.A., S-Y.L.; Methodology: I.R., S-Y.L., L.D.C., C.K., C.A.A; Software: I.R., C.A.A.; Writing – original draft: I.R., C.A.A.; Writing – review and edits: all authors.

## Code Availability

Code is available at the following link: https://github.com/anastassiou-team/NP_brain_stim_INVIVO

## Data Availability

Data analyzed in this manuscript are publicly available in: https://github.com/anastassiou-team/NP_brain_stim_INVIVO

## Acknowledgments

We thank the Allen Institute Animal Care, the Lab Animal Services teams for mouse husbandry and care; the Allen Institute Manufacturing and Process Engineering team for experimental hardware and software support; and the Imaging teams. We wish to thank the Allen Institute founder, Paul G. Allen, for his vision, encouragement, and support.

We gratefully acknowledge funding from the National Institute of Neurological Disorders and Stroke of the National Institutes of Health under Award Numbers R01NS120300 and RO1NS130126. The content is solely the responsibility of the authors and does not necessarily represent the official views of the National Institutes of Health.

## Online Methods

Experimental procedures closely followed those described in Claar et al. ^44^ and Russo et al. ^26^. A summary of these methods and details of procedures that differ are provided below.

### Mice

Mice were maintained in the Allen Institute animal facility and used in accordance with protocols approved by the Allen Institute’s Institutional Animal Care and Use Committee under protocol 2005. All experiments used C57BL/6J wild-type mice (N=17). Male and female wild-type C57BL/6J mice were purchased from Jackson Laboratories (JAX stock #000664) at postnatal day 28 and they were 9-28 weeks old at the time of all *in vivo* electrophysiological recordings. After surgery, all mice were single-housed and maintained on a reverse 12-h light cycle in a shared facility with room temperatures between 20 and 22 °C and humidity between 30 and 70%. All experiments were performed during the dark cycle. Mice had ad-libitum access to food and water.

### Surgical procedures and habituation

Each mouse went through the following order of procedures prior to the day of the experiment: 1) an initial sterile surgery to implant the reference and ground skull screws and a titanium headframe for head-fixed electrophysiological experiments *in vivo*; 2) five days of recovery time post-surgery; 3) at least three weeks of habituation to head-fixation; 4) and a second sterile surgery to perform small craniotomies to allow for insertion of the stimulating electrode and Neuropixels probes. Refer our protocols.io repository for details on surgical procedures, animal habituation and recording setup ^71^.

One to three hours prior to each surgery, pre-operative injections of dexamethasone (3-4 mg/kg, IM) and ceftriaxone (100-125 mg/kg, SC) were administered. Mice were deeply anesthetized with isoflurane (5% isoflurane induction, 1.5-2.5% maintenance) and placed in a stereotaxic frame. Vital signs were monitored, body temperature was maintained at 37.5 °C with a heating pad under the animal (TC-1000 temperature controller, CWE, Inc.), and ocular lubricant (I Drop, VetPLUS) was applied to maintain hydration of the eyes during anesthesia. Atropine (0.02-0.05 mg/kg, SC) and carprofen (5-10 mg/kg, SC) were administered at the start of the procedure. After the surgical procedure, mice received an injection of lactated Ringer’s solution (up to 1 mL, SC) and recovered on a heating pad. Animals received two days of analgesics and antibiotics post-surgery.

The initial surgery was performed on healthy mice that ranged in age from 5-20 weeks. Mice were deeply anesthetized prior to removing skin and exposing the skull. After leveling the skull, skull screws were implanted over the left and right cerebellum that functioned as reference and ground for the Neuropixels signals. White C&B Metabond (Parkell, Inc., Edgewood, New York) was then used to secure a custom titanium headframe to the skull. After five days of recovery, mice spent at least three weeks being habituated to handling and head-fixation.

Following habituation and up to one day before the recording, mice underwent the second surgical procedure. Under a microscope, the skull was exposed by drilling through the outer layer of Metabond. Up to two small craniotomies (less than 0.5 mm in diameter) were drilled to allow access to the brain regions of interest for the subsequent experiment. A small piece of artificial cerebrospinal fluid (ACSF)- soaked gel-foam sponge was positioned on top of each craniotomy and Kwik-Cast was used to seal it. A 3D-printed plastic well was also fixed to the existing Metabond around the craniotomies.

### *In-vivo* recording and stimulation

The day of the experiment, the mouse was placed on the running wheel and fixed to the headframe clamp with two set screws. Next, the thin layer of Kwik-Cast was removed to expose the craniotomies and abundant ACSF was added on top of the skull to prevent the exposed brain tissue from drying out. A 3D-printed cone was lowered to prevent the mouse’s tail from contacting the probes and a black curtain was lowered over the front of the rig, placing the mouse in complete darkness.

### Neuropixels recording

We performed recordings with two Neuropixels 1.0 probes^72^. These experiments used standard Neuropixels 1.0 probes configured to record from the 384 electrodes closest to the tip of the probe. The signals from each recording site were split in hardware into a spike band (30 kHz sampling rate, 500 Hz high-pass filter, 500x gain) and an LFP band (2.5 kHz sampling rate, 1,000 Hz low-pass filter, 250x gain) and data was acquired using the Open Ephys GUI ^73^. Each probe was connected to a PXI card inside a National Instruments chassis (Neuropixels 1.0).

The reference and ground connections on the Neuropixels probes were separated via two silver wires connected to two skull screws chronically implanted on top of the cerebellum area. All recordings were made using an external reference configuration (refer to our protocols.io repository for details ^71^). The mouse headframe and running wheel were shorted to the animal’s ground.

### Neuropixels insertion

During the acute electrophysiological experiment, up to two Neuropixels probes were inserted targeting visual area (VIS), hippocampal areas such as CA1, CA3 and dentate gyrus (DG) and other cortical areas anterior to VIS including somatosensory (SS) and retrosplenial (RSP).

The probe insertion process followed the steps detailed by Siegle, Jia et al. ^43^ and Claar et al. ^44^. Briefly, each probe was individually inserted into the brain using a 3-axis micromanipulator (New Scale Technologies, Victor, New York) at a rate of 200 μm per min to a depth of 3.5 mm or less in the brain. After the probes reached their targets, they were allowed to settle for 10-15 min before starting the experiment. Before insertion, all Neuropixels probes were coated with a fluorescent dye (Vybrant DiI/DiO/DiD, ThermoFisher Scientific, Waltham, Massachusetts) by repeatedly immersing them in a well filled with the dye and removing each probe slowly allowing the dye to dry on the surface.

### Behavioral data and synchronization

The angular position of the running wheel, and the synchronization signal for the electrical stimulus were acquired by a dedicated computer with a National Instruments card acquiring digital inputs at 100 kHz, which was considered the master clock. A 32-bit digital “barcode” was sent with an Arduino Uno (DEV-11021, SparkFun Electronics, Niwot, Colorado) every 30 s to synchronize all devices with the neural data from the Neuropixels probes. Details regarding the post-hoc data synchronization using the barcodes are described by Siegle, Jia et al.^43^. Videos of the right eye were acquired by a dedicated computer synchronized to the master clock at a sampling rate of 30Hz with the resolution of 640×480 pixels during the experimental sessions. The camera pointed to an infrared dichroic mirror placed in front of the right eye to allow the eye-tracking camera to operate without interference from the visual stimulus. The distance from the eye to mirror was ∼3cm and camera to mirror ∼9cm. To extract the pupillometry signal from the videos we developed a pipeline based on DeepLabCut (DLC) ^74^ and described in detail in Seyfourian et al. ^75^ with shared associated code.

### Cortical stimulation

Electrical stimulation was delivered through a custom bipolar platinum-iridium stereotrode (Microprobes for Life Science, Gaithersburg, Maryland) consisting of two parallel monopolar electrodes (50 kOhm impedance) with a vertical offset of 300 µm between the two tips. A stimulus generator (STG4002, Multichannel systems powered by Harvard Bioscience, Inc.) was set in current mode to generate and deliver continuous oscillatory stimulation at a fixed frequency and current intensity for 10 seconds-long sweeps. Each sweep followed by 10 seconds without stimulation was repeated 10 times constituting a block of stimulation at a specific set of stimulation parameters. Stimulation intensities applied were 1µA and 5µA at 8, 28 and 140 Hz for a total of six blocks of stimulation (three frequency and two intensities) and 60 sweeps (six blocks by ten sweeps) delivered to each animal. We recorded baseline conditions without stimulation for 5 minutes at the beginning and at the end of each block and at the beginning of each experiment. In two out of fourteen animals we delivered the same 60 sweeps arranged in the same six blocks but delivered in a randomized order.

The stimulation electrode was acutely inserted using a 3-axis micromanipulator, like the Neuropixels probes. We targeted three locations for stimulation: visual cortex (VIS), layer 5/6 (N=9 mice, 0.80±0.12 mm below the brain surface); somatosensory cortex (SS), layer 5/6 (N=4 mice, 0.94±0.05 mm below the brain surface); and secondary motor cortex (MOs), layer 5/6 (N=1 mouse, 1.22 mm below the brain surface), for a total number of 14 animals used for analyses (three were excluded based on signal quality control, see below for details). Before insertion, the stimulation electrode was coated with fluorescent dye, like the Neuropixels probes.

### Experimental timeline

The Neuropixels recordings were acute, lasting 2-3 hours. At the beginning, awake head-fixed mice (free to run or rest on the running wheel) were exposed to electrical stimuli after 5 min of baseline recording without stimulation. The resulting dataset allowed a direct comparison of the stimulation effects on neural activity across two behavioral states (quiet wakefulness and running).

### Probe removal and cleaning

Upon completion of the experiment, probes were retracted from the brain at a rate of 1 mm/s. The craniotomies were protected with a small piece of ACSF-soaked gel-foam sponge and a thin layer of Kwik-Cast before mice were removed from head fixation and returned to their home cages overnight. The probes and the stimulating electrode were immersed in a well of 1% Tergazyme for around 2-6 h to remove residual tissue and then rinsed in purified water.

### Ex vivo imaging and localization of electrodes

After the experiment, mice were deeply anesthetized (5% isoflurane) and perfused with 4% paraformaldehyde. The brains were preserved in 4% paraformaldehyde for 48 hours, rinsed with 1x PBS, and stored at 4 °C in PBS. The brains were then processed in one of two ways: brains were sliced into 100 µm coronal sections using a vibratome (Leica VT1000S) and imaged with a fluorescent microscope (Olympus VS110/120) at 10x magnification, or whole brains were imaged using serial two-photon tomography ^52,76^.

Images of the 100 µm coronal sections from serial two-photon tomography were aligned to the Allen Institute Common Coordinate Framework (CCFv3) following the process detailed by Oh et al. ^52^. Fluorescent tracks corresponding to the location of the Neuropixels probes and the stimulation electrode were manually identified in the aligned images. Because each CCFv3 coordinate corresponds to a unique brain region, the precision of brain region assignments is determined by the resolution of the CCFv3, which was 25 µm per pixel. For each Neuropixels probe the locations of major structural boundaries along the track was manually aligned with the physiology data ^43,77^. Then each recording channel along the Neuropixels probe (and associated neurons) was assigned to a unique CCFv3 structure. Other studies have reported better than 0.1 mm accuracy for electrode localization following similar methods ^77^.

### Data processing

#### Neuropixels LFP

All recordings were visually inspected by plotting the power spectra of the LFP signals across all channels. Recordings showing a clear 60 Hz peak in all LFP signals were excluded from further analysis. Based on this criterion, three animals were excluded, resulting in a final dataset of 14 animals. As described above, recordings were externally referenced to cerebellar skull screws, which allowed for the collection of reliable LFP signals without the need for additional re-referencing. The low stimulation intensities (1 µA and 5 µA) did not cause amplifier saturation; therefore, no stimulation artifact removal was required. Under these conditions, the applied currents effectively induced LFP oscillations at the targeted frequencies without introducing nonphysiological artifacts into the signals.

### Spike sorting

The raw action potential (AP) band data was pre-processed and spike-sorted using Kilosort 2.0 ^78^ as described in ^43^. High quality units were identified for further analysis using metrics per ^43^. Neurons were classified as regular spiking (RS) or fast spiking (FS)—putative pyramidal and inhibitory neurons, respectively—according to their spike waveform duration (RS: > 0.4 ms; FS: ≤ 0.4 ms) ^79–83^. To ensure that the applied stimulation did not alter spike waveforms, we applied an additional inclusion criterion based on the similarity between waveforms recorded at baseline (no stimulation) and during stimulation for each protocol. Spike waveforms were obtained by averaging all spikes from each unit within the respective time windows. The root mean square error (RMSE) between baseline and stimulation waveform templates was then calculated for each unit and protocol. Units with an RMSE greater than 0.10 were excluded from further analysis for that specific stimulation condition. This stringent criterion ensured that only stable, high-quality units were included, yielding a final total of 2,725 well-isolated units across 14 animals.

### Data Analysis

#### Population vector length (VL)

Units were included if they met predefined quality criteria, including a minimum spike count of ≥51 spikes per epoch and a circular fit error <0.1 during both baseline and stimulation periods. Brain areas were included only if at least 8 units satisfied these criteria. For spike-phase analysis, defined as the phase of the extracellular LFP at the time of each spike, the raw LFP traces were first bandpass filtered using a third-order Butterworth filter (scipy.signal.filtfilt) within ±20% of the frequency of interest. The instantaneous phase of the filtered LFPs was then calculated using the Hilbert transform. LFP signals were extracted from neighboring sites located 15–30 channels away from each unit’s peak channel to minimize contamination from spiking activity ^84^. Phases were expressed in degrees (0°–360°), with 90° corresponding to the LFP peak and 270° to the trough. The spike phase for each unit was defined as the instantaneous LFP phase at the spike time (see *Spike Sorting*, Methods).

Electrical stimulation (ES) entrainment was quantified using population vector analysis. For each unit, a mean vector length (VL; range 0–1) of spike phases was calculated to assess whether spikes were preferentially aligned to specific LFP phases. Spike-phase distributions were compared between baseline (pre-stimulation or control, 10 s) and stimulation (10 s) epochs for each trial. Each trial consisted of ten pre-stimulation and ten stimulation periods for a given stimulation frequency (8, 28, or 140 Hz) and amplitude (1 or 5 µA), yielding a total of 100 seconds of data per epoch. Increased entrainment was reflected by an inhomogeneous spike-phase distribution with a distinct preferred phase and a larger vector length (approaching 1), whereas weak or absent modulation resulted in a homogeneous distribution with a smaller vector length (approaching 0). The change in spike phase coupling during sES was quantified as ΔVL = VL_Stim − VL_Pre and tested for significance within each brain area using one-sample t-tests on the pairwise differences (scipy.stats.ttest_1samp). P-values were corrected across brain areas using the Benjamini–Hochberg false discovery rate (FDR) procedure (α = 0.05), with significance thresholds set at p < 0.01 and p < 0.001.

### Mean modulation ratio (MMR)

To quantify spike-phase entrainment we computed two complementary metrics for each unit: vector length (VL; see above) and the mean modulation ratio (MMR). VL, is the magnitude of vector summation of the circular spike-phase distribution and measures unimodal phase concentration. VL is well-defined for unimodal phase distributions but can yield artifactually low values when spikes cluster at two or more preferred phases, e.g. when out-of-phase unit vectors cancel out (for example, two units firing at 0° and 180° relative to the stimulation cycle produce VL ≈ 0 despite both units exhibiting strong coupling individually). To overcome this limitation we developed MMR, which quantifies any departure from phase uniformity regardless of the number of modes. For each unit satisfying the minimum spike-count criterion (≥51 spikes per epoch), spike times during both the pre-stimulation baseline and the stimulation epoch were converted to phase angles (0–360°) relative to the concurrent sES cycle. Each spike phase was represented as a Gaussian kernel (σ = 9°) placed on a 360-bin circular histogram (1° resolution), with circular wrapping at the 0°/360° boundary. The individual kernels were summed across all spikes to yield a continuous phase-occupancy distribution, which was then normalized to its peak value. MMR was defined as MMR = 1 - <*p*> where <*p*> denotes the mean of the peak-normalized distribution. Intuitively, a uniform distribution yields <*p*> close to 1 and MMR ≈ 0, whereas concentration of spikes at any number of preferred phases reduces <*p*> and increases MMR toward 1. The metric is therefore sensitive to unimodal, bimodal, and multimodal distributions and is well-suited for detecting multimodal (multi-peak) entrainment patterns observed during sES (Fig. S4). The change in phase coupling with stimulation was quantified as ΔMMR = MMR_Stim − MMR_Pre and tested for significance within each brain area using the same paired t-test and FDR correction procedure used for ΔVL. Despite their differing sensitivities, VL and MMR yielded highly concordant entrainment maps (Fig. 2 vs. Fig. S4), with MMR additionally detecting 20–40% more significantly entrained areas, predominantly at theta (8 Hz) and beta (28 Hz) frequencies where bimodal phase coupling was most prevalent.

### Spatial decay fitting and hierarchical regression analysis

To characterize the spatial decay of stimulation effects, the absolute change in vector length (|ΔVL|) was modeled as a function of distance from the stimulation site. Specifically, nonlinear least squares fitting of a double hyperbolic decay model was applied to unit-level |ΔVL| values using the model f(d) = c + 1/(d − d_0_)ⁿ,where d is distance in mm. Fits were performed using scipy.optimize.curve_fit with a robust cost function. This formulation captures the nonlinear decay of stimulation-induced effects with distance while allowing flexibility for near-field offsets and asymptotic baseline levels. To evaluate the contribution of multiple predictors, hierarchical ordinary least squares (OLS) regression was performed with all predictors z-scored prior to fitting. Nested models were compared using ΔR^2^ to quantify the incremental variance explained by the addition of each predictor set: Model 1 (distance only), Model 2 (distance + LFP amplitude at sES frequency), and Model 3 (full model including four anatomical connectivity metrics from the Allen Mouse Brain Connectivity Atlas). Significance of individual regression coefficients was assessed via two-sided t-tests from the OLS output.

### Clustering analysis

To identify distinct patterns of phase entrainment, clustering analyses were performed using Gaussian mixture models fitted to standardized features (VL during baseline and stimulation). The optimal number of clusters (k = 2–4) was determined by maximizing the silhouette score. Differences in entrainment across clusters were assessed using two-sided Mann–Whitney U tests for two-cluster solutions or Kruskal–Wallis tests for solutions with more than two clusters, with p-values FDR-corrected across brain areas. Associations between EAP waveform properties and cluster membership were tested using the same framework, with FDR correction applied across features within each area.

### Transient firing rate modulation

Transient changes in firing rate were quantified following a procedure adapted from Butovas & Schwarz (2003). Peristimulus time histograms (PSTHs) were computed as spike renewal densities (Abeles, 1982) with a temporal resolution of 1 ms to estimate firing rates over time around the stimulation onset. The baseline spontaneous firing rate (f_baseline_) was defined as the mean rate during the 10-second window preceding stimulation onset. Each PSTH was then smoothed by convolving it with a Gaussian kernel (kernel length: 50 bins).

Units were classified as excited if the firing rate increased to at least 1.1 × f_baseline_ within 10 seconds after stimulation onset. The duration of the excitatory response was defined as the time interval during which the firing rate remained above this threshold. Conversely, units were classified as inhibited if their firing rate fell below 0.95 × f_baseline_. The duration of the inhibitory response was measured from stimulation onset to the time point at which the firing rate returned above threshold.

To avoid detecting spurious fluctuations, a segment was considered a significant change only if it met all the following criteria: (1) a minimum absolute change in firing rate of 0.1 Hz; (2) a minimum duration of 100 ms; and (3) an onset occurring within 100 ms of stimulation onset. The strength of each response was defined as the absolute value of the Z-scored firing rate relative to baseline over the first 100 ms after stimulation onset. Change detection was mutually exclusive—if a unit exhibited an excitatory increase within the first 100 ms, it was classified as excited regardless of subsequent activity. To visualize population-level effects independent of response direction, we computed the absolute Z-scored firing rate for each unit. This was calculated over 100 ms following stimulation onset, relative to a 1-second pre stimulation baseline.

### Behavioral analysis

To evaluate how the animal’s behavioral state (quiet wakefulness vs. active state) influenced spike entrainment and transient firing rate responses, two complementary strategies were employed.

For the spike entrainment analysis, each spike from each unit was classified according to the animal’s behavioral state. A spike was labeled as *rest* if the mouse’s speed (measured from the wheel’s angular velocity) was below 0.3 cm/s during the ±0.1 s window around spike onset, and as *run* if the speed exceeded 0.3 cm/s. Given this additional classification and the requirement of a minimum of 51 spikes to compute the vector length (VL; see details above) for each behavioral condition and for both baseline (pre-stimulation) and stimulation periods, the number of units per brain area was reduced. However, thanks to the richness of the dataset, a relevant number of neurons could still be retained by pooling across subareas.

For the transient firing rate analysis, behavioral state classification was applied at the level of individual stimulation sweeps. A trial was classified as *rest* if the mouse’s speed was below 0.1 cm/s within the −1 s to +0.5 s window relative to stimulation onset, and as *run* if the speed exceeded 0.1 cm/s during this same period. Because transient firing rate effects were quantified from peristimulus time histograms (PSTHs), each behavioral condition required a minimum of three valid trials to be included in the analysis.

To assess whether stimulation onsets and offsets induced changes in behavior, we computed stimulation-triggered averages of both the pupillometry and wheel-speed signals. For each event, we extracted signal segments from −2 to +2 seconds around the onset or offset and averaged these single-trial sweeps to obtain the corresponding event-triggered responses.

### Statistics

Statistical analyses were performed in Python using SciPy (v1.15.3) and statsmodels (v0.14.5). Within-area comparisons of firing rate changes between baseline and stimulation (Fig. 4b) were assessed using the same paired t-test approach used for ΔVL and MMR. In all figures, the convention is *p < 0.05, **p < 0.01, ***p < 0.001, and ****p < 0.0001.

## References

1 Cerletti, U. & Bini, L. Electroshock (*)(dagger). Int Rev Psychiatry 30, 153–154, doi:10.1080/09540261.2018.1436662 (2018).

2 Mayberg, H. S. et al. Deep brain stimulation for treatment-resistant depression. Neuron 45, 651–660, doi:10.1016/j.neuron.2005.02.014 (2005).

3 Kringelbach, M. L., Jenkinson, N., Owen, S. L. & Aziz, T. Z. Translational principles of deep brain stimulation. Nat Rev Neurosci 8, 623–635, doi:10.1038/nrn2196 (2007).

4 Hogg, E. et al. Sustained quality-of-life improvements over 10 years after deep brain stimulation for dystonia. Mov Disord 33, 1160–1167, doi:10.1002/mds.27426 (2018).

5 Rektorova, I., Pupikova, M., Fleury, L., Brabenec, L. & Hummel, F. C. Non-invasive brain stimulation: current and future applications in neurology. Nat Rev Neurol 21, 669–686, doi:10.1038/s41582-025-01137-z (2025).

6 Wostmann, M., Vosskuhl, J., Obleser, J. & Herrmann, C. S. Opposite effects of lateralised transcranial alpha versus gamma stimulation on auditory spatial attention. Brain Stimul 11, 752–758, doi:10.1016/j.brs.2018.04.006 (2018).

7 Helfrich, R. F. et al. Selective modulation of interhemispheric functional connectivity by HD-tACS shapes perception. PLoS Biol 12, e1002031, doi:10.1371/journal.pbio.1002031 (2014).

8 Alekseichuk, I., Turi, Z., Veit, S. & Paulus, W. Model-driven neuromodulation of the right posterior region promotes encoding of long-term memories. Brain Stimul 13, 474–483, doi:10.1016/j.brs.2019.12.019 (2020).

9 Takeuchi, Y. & Berenyi, A. Oscillotherapeutics - Time-targeted interventions in epilepsy and beyond. Neurosci Res 152, 87–107, doi:10.1016/j.neures.2020.01.002 (2020).

10 Stoney, S. D., Jr., Thompson, W. D. & Asanuma, H. Excitation of pyramidal tract cells by intracortical microstimulation: effective extent of stimulating current. J Neurophysiol 31, 659–669, doi:10.1152/jn.1968.31.5.659 (1968).

11 Nowak, L. G. & Bullier, J. Axons, but not cell bodies, are activated by electrical stimulation in cortical gray matter. I. Evidence from chronaxie measurements. Exp Brain Res 118, 477–488, doi:10.1007/s002210050304 (1998).

12 Tehovnik, E. J., Tolias, A. S., Sultan, F., Slocum, W. M. & Logothetis, N. K. Direct and indirect activation of cortical neurons by electrical microstimulation. J Neurophysiol 96, 512–521, doi:10.1152/jn.00126.2006 (2006).

13 Vieira, P. G., Krause, M. R., Laamerad, P. & Pack, C. C. Brain stimulation preferentially influences long-range projections. Sci Adv 11, eadx2106, doi:10.1126/sciadv.adx2106 (2025).

14 Radman, T., Ramos, R. L., Brumberg, J. C. & Bikson, M. Role of cortical cell type and morphology in subthreshold and suprathreshold uniform electric field stimulation in vitro. Brain Stimul 2, 215–228, 228 e211-213, doi:10.1016/j.brs.2009.03.007 (2009).

15 Ye, H. & Steiger, A. Neuron matters: electric activation of neuronal tissue is dependent on the interaction between the neuron and the electric field. J Neuroeng Rehabil 12, 65, doi:10.1186/s12984-015-0061-1 (2015).

16 Bikson, M. et al. Effects of uniform extracellular DC electric fields on excitability in rat hippocampal slices in vitro. J Physiol 557, 175–190, doi:10.1113/jphysiol.2003.055772 (2004).

17 Tolias, A. S. et al. Mapping cortical activity elicited with electrical microstimulation using FMRI in the macaque. Neuron 48, 901–911, doi:10.1016/j.neuron.2005.11.034 (2005).

18 Richardson, A. G., McIntyre, C. C. & Grill, W. M. Modelling the effects of electric fields on nerve fibres: influence of the myelin sheath. Med Biol Eng Comput 38, 438–446, doi:10.1007/BF02345014 (2000).

19 McIntyre, C. C. & Grill, W. M. Selective microstimulation of central nervous system neurons. Ann Biomed Eng 28, 219–233, doi:10.1114/1.262 (2000).

20 McIntyre, C. C. & Grill, W. M. Extracellular stimulation of central neurons: influence of stimulus waveform and frequency on neuronal output. J Neurophysiol 88, 1592–1604, doi:10.1152/jn.2002.88.4.1592 (2002).

21 Kumaravelu, K., Sombeck, J., Miller, L. E., Bensmaia, S. J. & Grill, W. M. Stoney vs. Histed: Quantifying the spatial effects of intracortical microstimulation. Brain Stimul 15, 141–151, doi:10.1016/j.brs.2021.11.015 (2022).

22 Histed, M. H., Bonin, V. & Reid, R. C. Direct activation of sparse, distributed populations of cortical neurons by electrical microstimulation. Neuron 63, 508–522, doi:10.1016/j.neuron.2009.07.016 (2009).

23 Butovas, S. & Schwarz, C. Spatiotemporal effects of microstimulation in rat neocortex: a parametric study using multielectrode recordings. J Neurophysiol 90, 3024–3039, doi:10.1152/jn.00245.2003 (2003).

24 Logothetis, N. K. et al. The effects of electrical microstimulation on cortical signal propagation. Nat Neurosci 13, 1283–1291, doi:10.1038/nn.2631 (2010).

25 Yun, R., Mishler, J. H., Perlmutter, S. I., Rao, R. P. N. & Fetz, E. E. Responses of Cortical Neurons to Intracortical Microstimulation in Awake Primates. eNeuro 10, doi:10.1523/ENEURO.0336-22.2023 (2023).

26 Russo, S. et al. Thalamic feedback shapes brain responses evoked by cortical stimulation in mice and humans. Nat Commun 16, 3627, doi:10.1038/s41467-025-58717-2 (2025).

27 Ozen, S. et al. Transcranial electric stimulation entrains cortical neuronal populations in rats. J Neurosci 30, 11476–11485, doi:10.1523/JNEUROSCI.5252-09.2010 (2010).

28 Krause, M. R., Vieira, P. G., Csorba, B. A., Pilly, P. K. & Pack, C. C. Transcranial alternating current stimulation entrains single-neuron activity in the primate brain. Proc Natl Acad Sci U S A 116, 5747–5755, doi:10.1073/pnas.1815958116 (2019).

29 Krause, M. R., Vieira, P. G. & Pack, C. C. Transcranial electrical stimulation: How can a simple conductor orchestrate complex brain activity? PLoS Biol 21, e3001973, doi:10.1371/journal.pbio.3001973 (2023).

30 Krause, M. R., Vieira, P. G., Thivierge, J. P. & Pack, C. C. Brain stimulation competes with ongoing oscillations for control of spike timing in the primate brain. PLoS Biol 20, e3001650, doi:10.1371/journal.pbio.3001650 (2022).

31 Johnson, L. et al. Dose-dependent effects of transcranial alternating current stimulation on spike timing in awake nonhuman primates. Sci Adv 6, doi:10.1126/sciadv.aaz2747 (2020).

32 Ahn, S. et al. Targeting reduced neural oscillations in patients with schizophrenia by transcranial alternating current stimulation. Neuroimage 186, 126–136, doi:10.1016/j.neuroimage.2018.10.056 (2019).

33 Alexander, M. L. et al. Double-blind, randomized pilot clinical trial targeting alpha oscillations with transcranial alternating current stimulation (tACS) for the treatment of major depressive disorder (MDD). Transl Psychiatry 9, 106, doi:10.1038/s41398-019-0439-0 (2019).

34 Hyde, J. et al. Efficacy of neurostimulation across mental disorders: systematic review and meta-analysis of 208 randomized controlled trials. Mol Psychiatry 27, 2709–2719, doi:10.1038/s41380-022-01524-8 (2022).

35 Holtzheimer, P. E. et al. Subcallosal cingulate deep brain stimulation for treatment-resistant depression: a multisite, randomised, sham-controlled trial. Lancet Psychiatry 4, 839–849, doi:10.1016/S2215-0366(17)30371-1 (2017).

36 Heroux, M. E., Loo, C. K., Taylor, J. L. & Gandevia, S. C. Questionable science and reproducibility in electrical brain stimulation research. PLoS One 12, e0175635, doi:10.1371/journal.pone.0175635 (2017).

37 Gradinaru, V., Mogri, M., Thompson, K. R., Henderson, J. M. & Deisseroth, K. Optical deconstruction of parkinsonian neural circuitry. Science 324, 354–359, doi:10.1126/science.1167093 (2009).

38 Borchers, S., Himmelbach, M., Logothetis, N. & Karnath, H. O. Direct electrical stimulation of human cortex - the gold standard for mapping brain functions? Nat Rev Neurosci 13, 63–70, doi:10.1038/nrn3140 (2011).

39 Bradley, C., Nydam, A. S., Dux, P. E. & Mattingley, J. B. State-dependent effects of neural stimulation on brain function and cognition. Nat Rev Neurosci 23, 459–475, doi:10.1038/s41583-022-00598-1 (2022).

40 Voroslakos, M. et al. Direct effects of transcranial electric stimulation on brain circuits in rats and humans. Nat Commun 9, 483, doi:10.1038/s41467-018-02928-3 (2018).

41 Anastassiou, C. A., Perin, R., Markram, H. & Koch, C. Ephaptic coupling of cortical neurons. Nat Neurosci 14, 217–223, doi:10.1038/nn.2727 (2011).

42 Lee, S. Y. et al. Cell-class-specific electric field entrainment of neural activity. Neuron 112, 2659–2660, doi:10.1016/j.neuron.2024.07.011 (2024).

43 Siegle, J. H. et al. Survey of spiking in the mouse visual system reveals functional hierarchy. Nature 592, 86–92, doi:10.1038/s41586-020-03171-x (2021).

44 Claar, L. D. et al. Cortico-thalamo-cortical interactions modulate electrically evoked EEG responses in mice. Elife 12, doi:10.7554/eLife.84630 (2023).

45 Harris, J. A. et al. Hierarchical organization of cortical and thalamic connectivity. Nature 575, 195–202, doi:10.1038/s41586-019-1716-z (2019).

46 Knox, J. E. et al. High-resolution data-driven model of the mouse connectome. Netw Neurosci 3, 217–236, doi:10.1162/netn_a_00066 (2019).

47 Anastassiou, C. A., Montgomery, S. M., Barahona, M., Buzsaki, G. & Koch, C. The effect of spatially inhomogeneous extracellular electric fields on neurons. J Neurosci 30, 1925–1936, doi:10.1523/JNEUROSCI.3635-09.2010 (2010).

48 Olsen, S. R., Bortone, D. S., Adesnik, H. & Scanziani, M. Gain control by layer six in cortical circuits of vision. Nature 483, 47–52, doi:10.1038/nature10835 (2012).

49 Busse, L. et al. Sensation during Active Behaviors. J Neurosci 37, 10826–10834, doi:10.1523/JNEUROSCI.1828-17.2017 (2017).

50 Niell, C. M. & Stryker, M. P. Modulation of visual responses by behavioral state in mouse visual cortex. Neuron 65, 472–479, doi:10.1016/j.neuron.2010.01.033 (2010).

51 Dadarlat, M. C. & Stryker, M. P. Locomotion Enhances Neural Encoding of Visual Stimuli in Mouse V1. J Neurosci 37, 3764–3775, doi:10.1523/JNEUROSCI.2728-16.2017 (2017).

52 Oh, S. W. et al. A mesoscale connectome of the mouse brain. Nature 508, 207–214, doi:10.1038/nature13186 (2014).

53 Fox, K. C. R. et al. Intrinsic network architecture predicts the effects elicited by intracranial electrical stimulation of the human brain. Nat Hum Behav 4, 1039–1052, doi:10.1038/s41562-020-0910-1 (2020).

54 Lyu, D., Stiger, J. R., Lusk, Z., Buch, V. & Parvizi, J. Mapping human thalamocortical connectivity with electrical stimulation and recording. Nat Neurosci 28, 1797–1809, doi:10.1038/s41593-025-02009-x (2025).

55 Keller, C. J. et al. Intrinsic functional architecture predicts electrically evoked responses in the human brain. Proc Natl Acad Sci U S A 108, 10308–10313, doi:10.1073/pnas.1019750108 (2011).

56 Matsumoto, R., Kunieda, T. & Nair, D. Single pulse electrical stimulation to probe functional and pathological connectivity in epilepsy. Seizure 44, 27–36, doi:10.1016/j.seizure.2016.11.003 (2017).

57 Matsumoto, R. et al. Functional connectivity in human cortical motor system: a cortico-cortical evoked potential study. Brain 130, 181–197, doi:10.1093/brain/awl257 (2007).

58 Matsumoto, R. et al. Functional connectivity in the human language system: a cortico-cortical evoked potential study. Brain 127, 2316–2330, doi:10.1093/brain/awh246 (2004).

59 Frohlich, F. & McCormick, D. A. Endogenous electric fields may guide neocortical network activity. Neuron 67, 129–143, doi:10.1016/j.neuron.2010.06.005 (2010).

60 Overstreet, C. K., Klein, J. D. & Helms Tillery, S. I. Computational modeling of direct neuronal recruitment during intracortical microstimulation in somatosensory cortex. J Neural Eng 10, 066016, doi:10.1088/1741-2560/10/6/066016 (2013).

61 Buzsaki, G. Neural syntax: cell assemblies, synapsembles, and readers. Neuron 68, 362–385, doi:10.1016/j.neuron.2010.09.023 (2010).

62 McIntyre, C. C., Grill, W. M., Sherman, D. L. & Thakor, N. V. Cellular effects of deep brain stimulation: model-based analysis of activation and inhibition. J Neurophysiol 91, 1457–1469, doi:10.1152/jn.00989.2003 (2004).

63 McGinley, M. J. et al. Waking State: Rapid Variations Modulate Neural and Behavioral Responses. Neuron 87, 1143–1161, doi:10.1016/j.neuron.2015.09.012 (2015).

64 Hu, H., Gan, J. & Jonas, P. Interneurons. Fast-spiking, parvalbumin(+) GABAergic interneurons: from cellular design to microcircuit function. Science 345, 1255263, doi:10.1126/science.1255263 (2014).

65 Cardin, J. A. et al. Driving fast-spiking cells induces gamma rhythm and controls sensory responses. Nature 459, 663–667, doi:10.1038/nature08002 (2009).

66 Csicsvari, J., Hirase, H., Czurko, A. & Buzsaki, G. Reliability and state dependence of pyramidal cell-interneuron synapses in the hippocampus: an ensemble approach in the behaving rat. Neuron 21, 179–189, doi:10.1016/s0896-6273(00)80525-5 (1998).

67 Uhlhaas, P. J. & Singer, W. Abnormal neural oscillations and synchrony in schizophrenia. Nat Rev Neurosci 11, 100–113, doi:10.1038/nrn2774 (2010).

68 Berenyi, A., Belluscio, M., Mao, D. & Buzsaki, G. Closed-loop control of epilepsy by transcranial electrical stimulation. Science 337, 735–737, doi:10.1126/science.1223154 (2012).

69 Rembado, I., Zanos, S. & Fetz, E. E. Cycle-Triggered Cortical Stimulation during Slow Wave Sleep Facilitates Learning a BMI Task: A Case Report in a Non-Human Primate. Front Behav Neurosci 11, 59, doi:10.3389/fnbeh.2017.00059 (2017).

70 Zanos, S., Rembado, I., Chen, D. & Fetz, E. E. Phase-Locked Stimulation during Cortical Beta Oscillations Produces Bidirectional Synaptic Plasticity in Awake Monkeys. Curr Biol 28, 2515–2526 e2514, doi:10.1016/j.cub.2018.07.009 (2018).

71 Marks, L. C., Rembado, I. & Claar, L. D. . Simultaneous Recording of Neuropixels and EEG in Head-Fixed Mice. protocols.io 10.17504/protocols.io.14egnSq1Cl5d/v1.

72 Jun, J. J. et al. Fully integrated silicon probes for high-density recording of neural activity. Nature 551, 232–236, doi:10.1038/nature24636 (2017).

73 Siegle, J. H. et al. Open Ephys: an open-source, plugin-based platform for multichannel electrophysiology. J Neural Eng 14, 045003, doi:10.1088/1741-2552/aa5eea (2017).

74 Mathis, A. et al. DeepLabCut: markerless pose estimation of user-defined body parts with deep learning. Nat Neurosci 21, 1281–1289, doi:10.1038/s41593-018-0209-y (2018).

75 Seyfourian P., M. L. C., Claar L. D., Nahas Y., Keating M., Koch C., Rembado I. Pupil-DLC: an open-source deep learning pipeline for scalable, markerless tracking of pupil dynamics across conscious and unconscious states. biorxiv, 10.64898/2026.01.18.700183 (2026).

76 Ragan, T. et al. Serial two-photon tomography for automated ex vivo mouse brain imaging. Nat Methods 9, 255–258, doi:10.1038/nmeth.1854 (2012).

77 Liu, L. D. et al. Accurate Localization of Linear Probe Electrode Arrays across Multiple Brains. eNeuro 8, doi:10.1523/ENEURO.0241-21.2021 (2021).

78 Stringer, C. et al. Spontaneous behaviors drive multidimensional, brainwide activity. Science 364, 255, doi:10.1126/science.aav7893 (2019).

79 Bartho, P. et al. Characterization of neocortical principal cells and interneurons by network interactions and extracellular features. J Neurophysiol 92, 600–608, doi:10.1152/jn.01170.2003 (2004).

80 Bruno, R. M. & Simons, D. J. Feedforward mechanisms of excitatory and inhibitory cortical receptive fields. J Neurosci 22, 10966–10975, doi:10.1523/JNEUROSCI.22-24-10966.2002 (2002).

81 Bortone, D. S., Olsen, S. R. & Scanziani, M. Translaminar inhibitory cells recruited by layer 6 corticothalamic neurons suppress visual cortex. Neuron 82, 474–485, doi:10.1016/j.neuron.2014.02.021 (2014).

82 Niell, C. M. & Stryker, M. P. Highly selective receptive fields in mouse visual cortex. J Neurosci 28, 7520–7536, doi:10.1523/JNEUROSCI.0623-08.2008 (2008).

83 Huo, Y., Chen, H. & Guo, Z. V. Mapping Functional Connectivity from the Dorsal Cortex to the Thalamus. Neuron 107, 1080–1094 e1085, doi:10.1016/j.neuron.2020.06.038 (2020).

84 Liu, A. A. et al. A consensus statement on detection of hippocampal sharp wave ripples and differentiation from other fast oscillations. Nat Commun 13, 6000, doi:10.1038/s41467-022-33536-x (2022).

