## Supplementary material for "Selective gating of neural modulation through frequency- and behavior-dependent modes during cortical electrical stimulation": SupplMat

### SUPPLEMENTAL MATERIAL

**Supplemental figures**

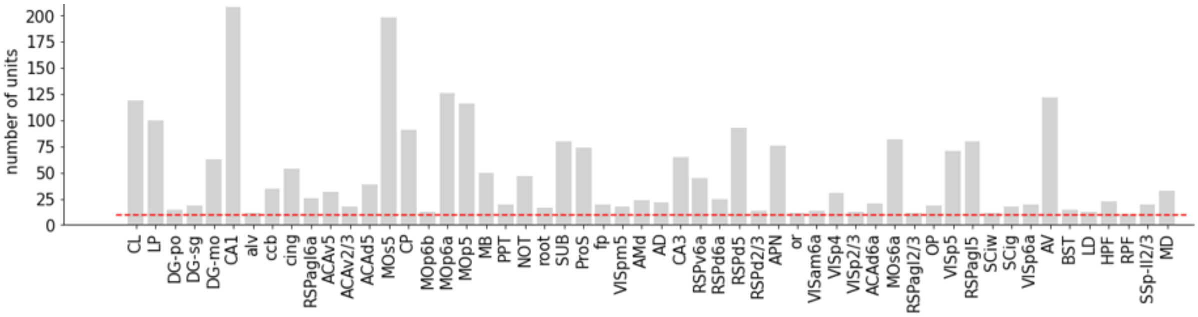


**Figure S1. Number of single units per brain area.** Bar plot showing the number of recorded single units assayed in each brain area across all subjects (*n*=14). Only areas with a minimum of 8 units (red dashed line) were included in further analyses.

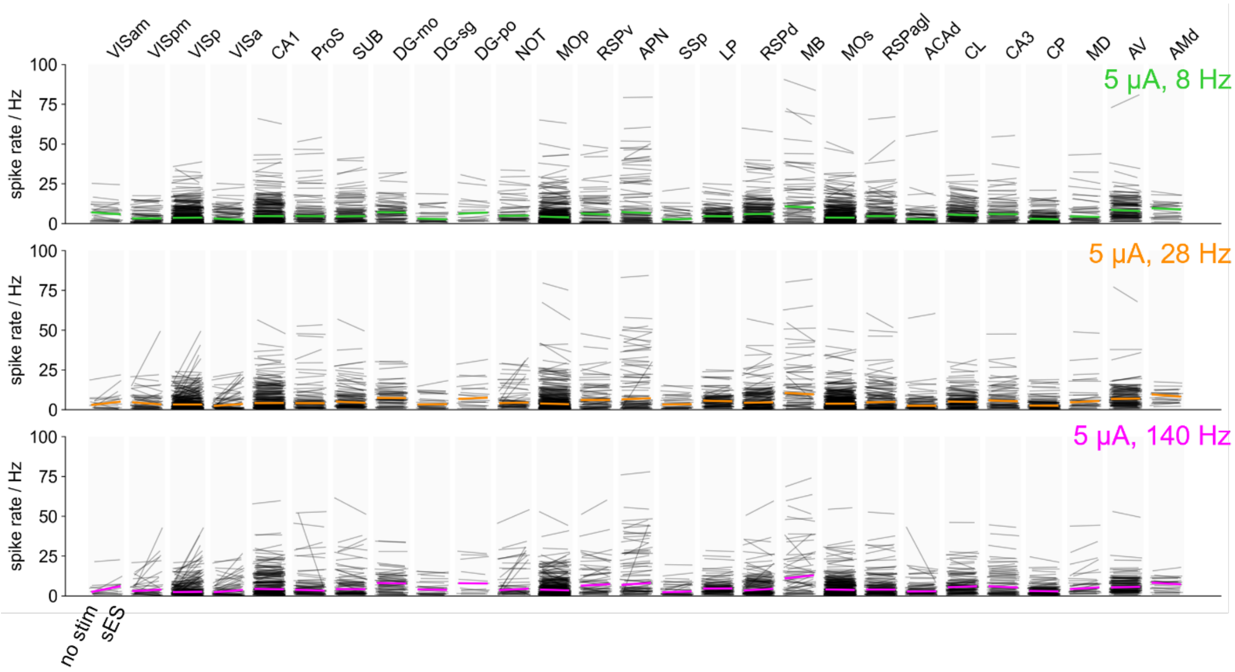


**Figure S2. Unit firing rate estimated during the entire sinusoidal electrical stimulation duration (sES) remains unchanged across brain areas.** Spike rates of individual units (gray lines) during the pre-stimulus baseline and the full 10-second stimulation epoch, shown for each brain area at 5 µA across all stimulation frequencies (8 Hz, green; 28 Hz, orange; 140 Hz, magenta). Colored horizontal lines indicate the median spike rate per stimulation protocol. No systematic firing rate changes were observed relative to baseline. Statistical testing: Mann-Whitney with Benjamini-Hochberg multiple comparison correction.

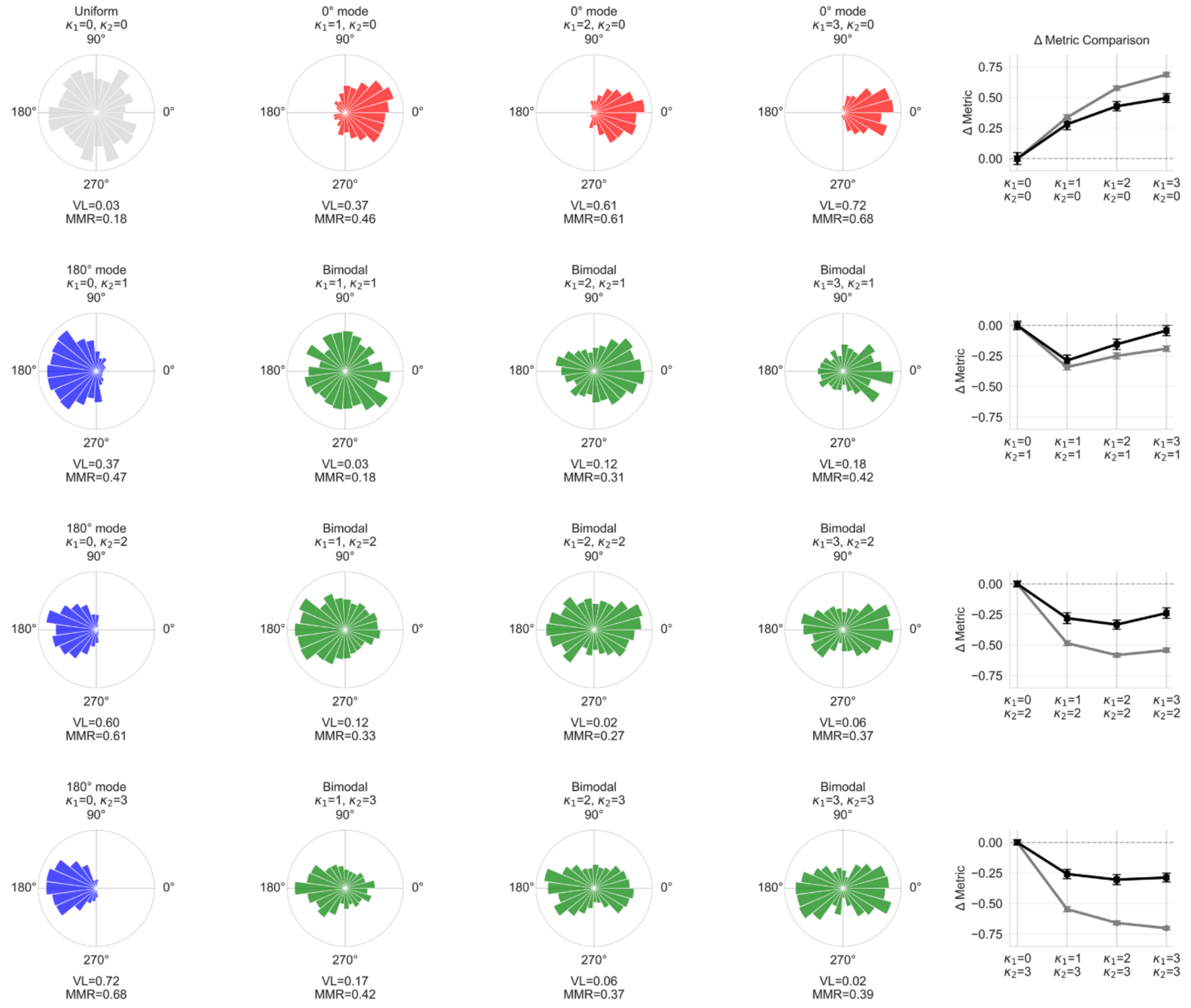


**Figure S3. Introducing the mean modulation ratio (MMR) and how it compares to VL in quantifying spike-phase entrainment for complex (multimodal) coupling profiles.** MMR is a more effective metric of spike phase coupling strength for multimodal entrainment compared to the population vector. Left column: (top) uniform distribution of 1,000 phases; (three next rose plots): unimodal phase distributions of increasing kappa (kappa = 1, 2, 3, respectively). (Top row): starting from a uniform phase distribution, unimodal phase distributions of increasing kappa (kappa = 1, 2, 3, respectively) exactly like in the left column but with different center phase (blue distributions: center phase is 180°; red distributions: center phase is 0°). The green distributions are sampled from the corresponding blue and red distributions (500 phases of each) generating bimodal distributions where each mode represents a different kappa and two center phases at 0° and 180°. (Right column) We define ΔVL= VL-VL (unimodal) and ΔMMR= MMR-MMR (unimodal) and show how they vary for the distributions in each row (circles: median; error bar: standard error of the mean; n=50 realizations of MMR and VL for each rose plot). We show that for bimodal distributions with pronounced modes, the VL is much more affected compared to MMR. Specifically, even very pronounced bimodal distributions result in large deviations of ΔVL (i.e., effectively a small VL) suggesting lack of phase locking. In contrast. MMR is much less affected by the presence of bimodal distributions and deviates less from its original value (i.e. coupling to a unimodal distribution). This renders the MMR a more appropriate metric to estimate phase locking effects in the presence of multiple modes.

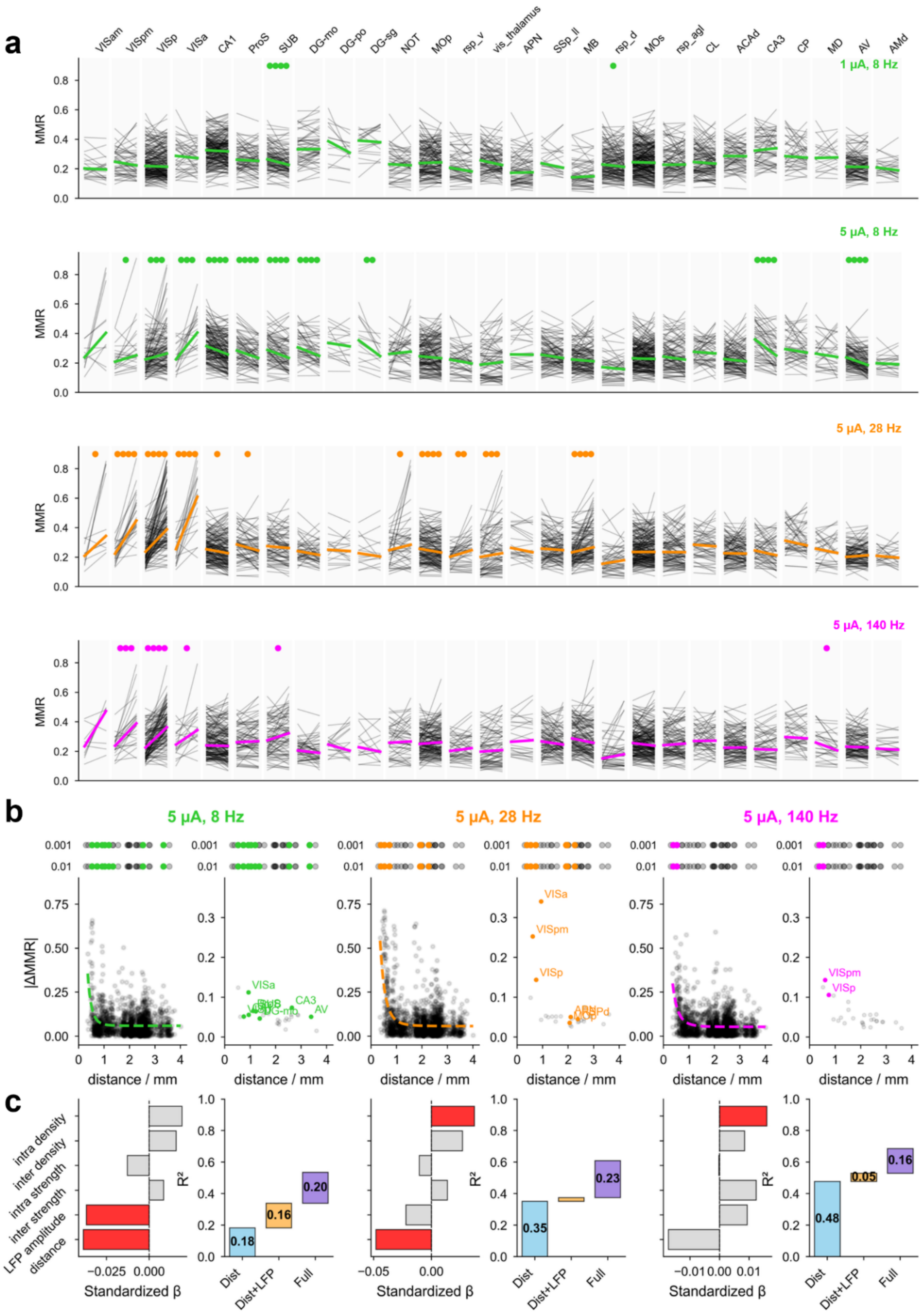
a
VISam VISpm VISp VISa CA1 ProS SUB DG-mo DG-po DG-sg NOT MOp rsp_v vis_thalamus APN SSp_ll MB rsp_d MOs rsp_agl CL ACAd CA3 CP MD AV AMd
1 µA, 8 Hz
5 µA, 8 Hz
5 µA, 28 Hz
5 µA, 140 Hz
MMR
b
5 µA, 8 Hz
5 µA, 28 Hz
5 µA, 140 Hz
|ΔMMR|
distance / mm
VISa
CA3
AV
VISpm
VISp
c
intra density
inter density
intra strength
inter strength
LFP amplitude
distance
Standardized β
R²
Dist
Dist+LFP
Full
0.18
0.16
0.20
0.35
0.23
0.48
0.05
0.16

64

**Figure S4. Spike-phase entrainment measured by mean modulation ratio (MMR) confirms results obtained with vector length (VL).** a) Spike-phase entrainment measured by MMR compared between baseline and sES across brain areas for four sES conditions (top to bottom: 1 µA 8 Hz, 5 µA 8 Hz, 5 µA 28 Hz, 5 µA 140 Hz; lines connect the same unit between baseline and sES). Statistical significance of MMR difference shown above each area (circle color corresponds to sES frequency; number of circles indicates significance level: ●●●● $p \leq 0.0001$; ●●● $p \leq 0.001$; ●● $p \leq 0.01$; ● $p \leq 0.05$; paired t-test). Brain areas ordered by increasing distance from sES location (left to right). b) Spatial decay of $|\Delta MMR|$ for 5 µA at 8, 28, and 140 Hz. Unit-level (left) and area-level (right) absolute change in MMR plotted against distance from stimulation site. Colored circles indicate significantly entrained areas ($p < 0.001$); gray circles indicate non-significant areas. c) Hierarchical regression analysis quantifying contributions of distance, LFP amplitude, and anatomical connectivity metrics to MMR-based entrainment strength across brain areas. Red bars: significant predictors ($p < 0.05$); gray bars: non-significant. Right panels: cumulative $R^2$ from hierarchical regression (blue: distance alone; orange: distance + LFP amplitude; purple: full model). Numbers indicate $\Delta R^2$ per model step.

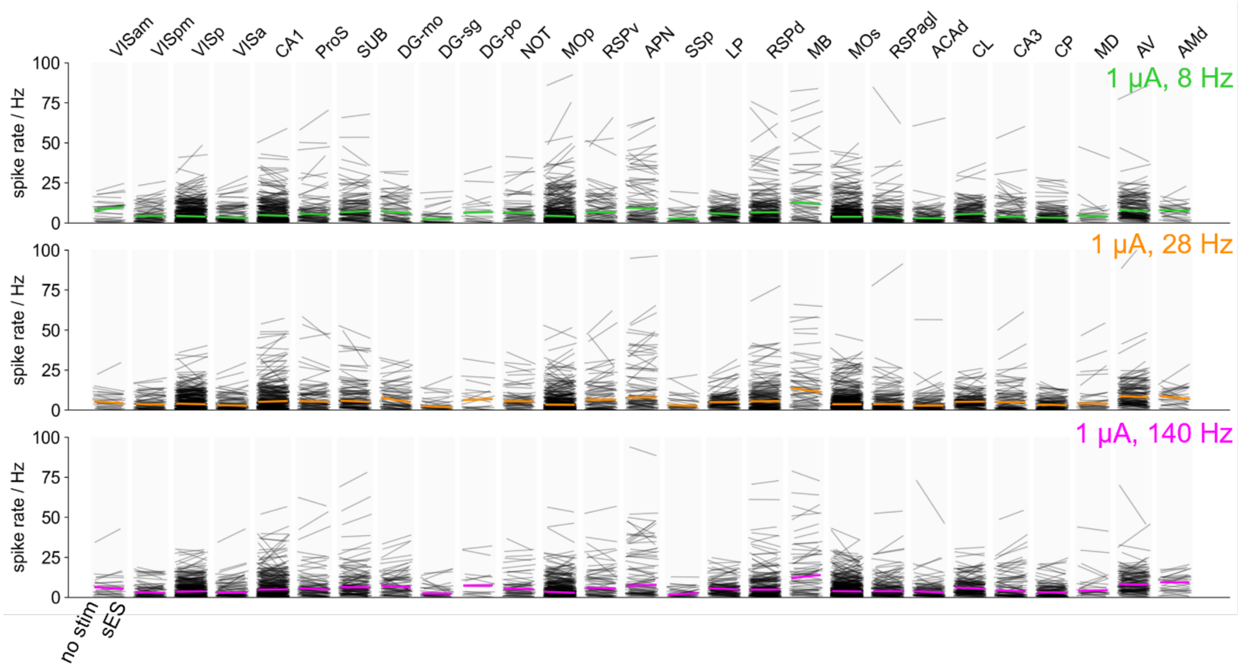


**Figure S5. Transient firing rate changes for 1 µA sES.** Transient spike rate modulation at stimulation onset for 1 µA across all stimulation frequencies and brain areas. No systematic firing rate changes were observed relative to baseline. Statistical testing: Mann-Whitney U-testing with Benjamini-Hochberg multiple comparison correction.

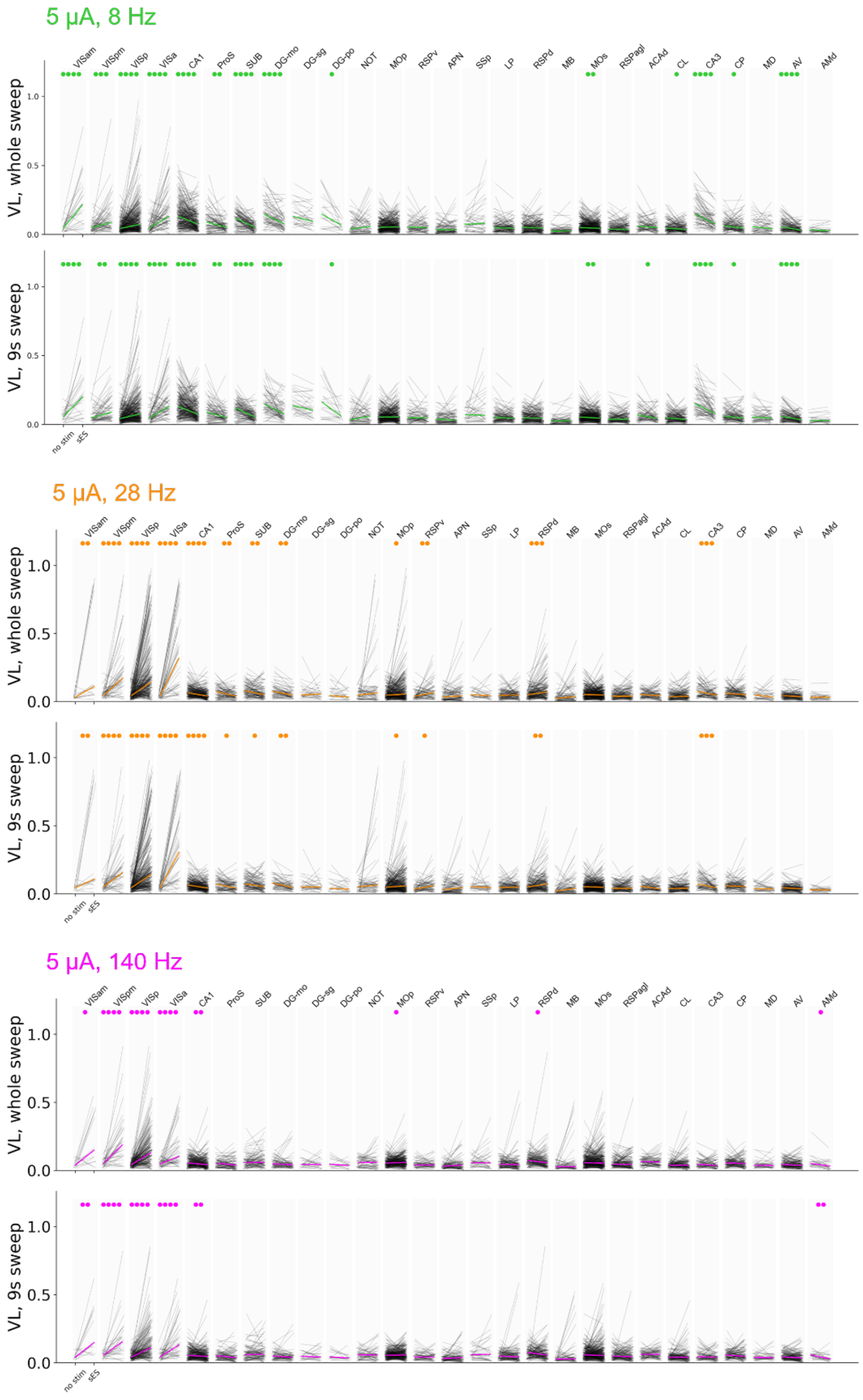
5 µA, 8 Hz
VL, whole sweep
VL, 9s sweep
VISam VISpm VISp VISa CA1 ProS SUB DG-mo DG-sg DG-po NOT MOp RSPv APN SSp LP RSPd MB MOs RSPagl ACAd CL CA3 CP MD AV AMd
no stim
sES
5 µA, 28 Hz
VL, whole sweep
VL, 9s sweep
VISam VISpm VISp VISa CA1 ProS SUB DG-mo DG-sg DG-po NOT MOp RSPv APN SSp LP RSPd MB MOs RSPagl ACAd CL CA3 CP MD AV AMd
no stim
sES
5 µA, 140 Hz
VL, whole sweep
VL, 9s sweep
VISam VISpm VISp VISa CA1 ProS SUB DG-mo DG-sg DG-po NOT MOp RSPv APN SSp LP RSPd MB MOs RSPagl ACAd CL CA3 CP MD AV AMd
no stim
sES

**Figure S6. Brain-wide spike-phase entrainment does not depend on transient firing rate effects.** Vector length (VL) across brain areas for all stimulation protocols at 5 µA (8 Hz, green; 28 Hz, orange; 140 Hz, magenta). For each frequency, the top panel shows VL calculated over the full 10-second stimulation epoch, while the bottom panel excludes the first second of stimulation to remove transient onset effects. Gray lines indicate individual units; colored lines indicate the median VL per area. Statistical significance shown above each area (●●●● $p \leq 0.0001$; ●●● $p \leq 0.001$; ●● $p \leq 0.01$; ● $p \leq 0.05$; Mann-Whitney test with Benjamini-Hochberg correction for multiple comparisons). The consistency of entrainment patterns between top and bottom panels confirms that brain-wide phase entrainment is not driven by transient spike-rate modulation at stimulation onset.

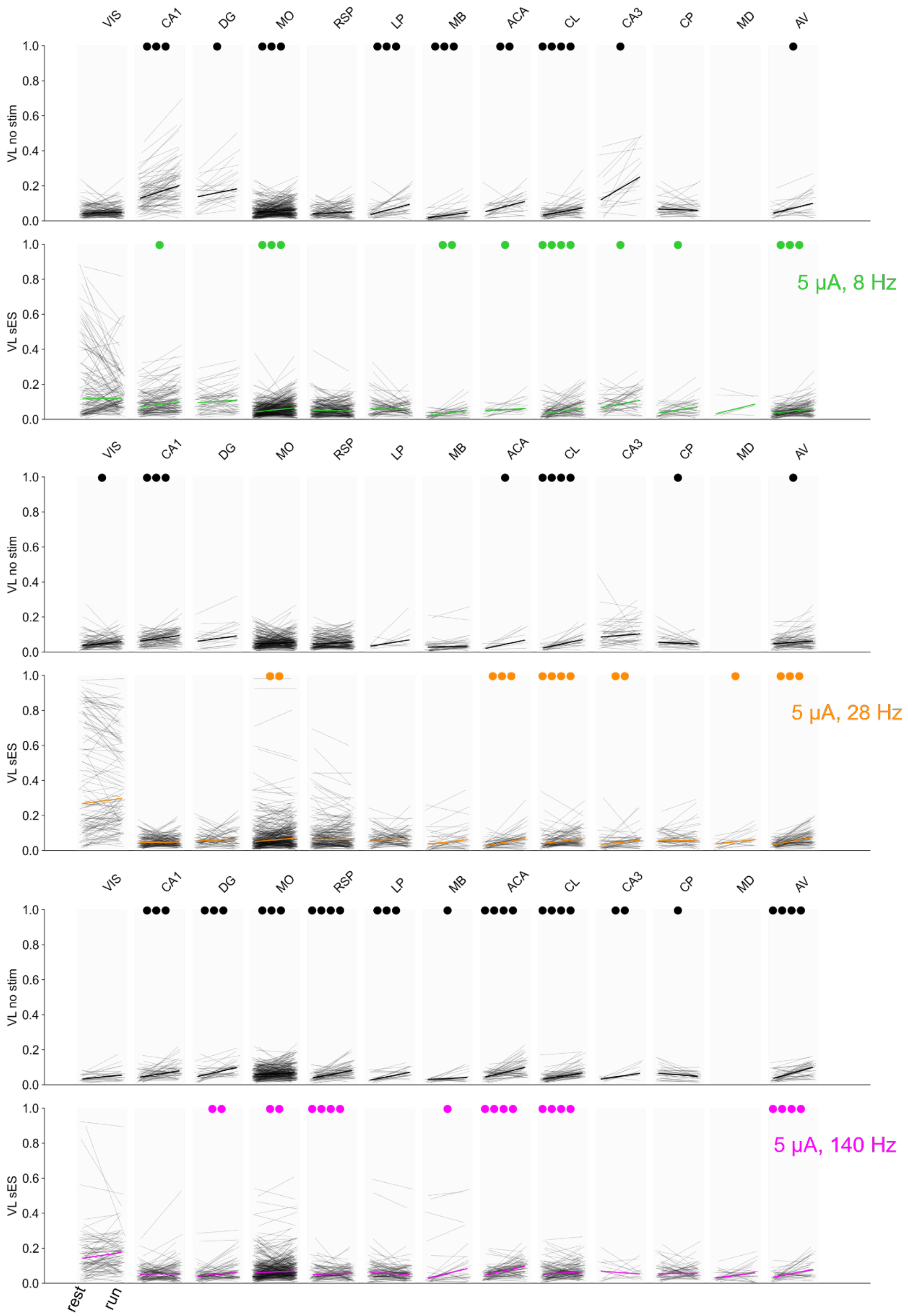
VIS
CA1
DG
MO
RSP
LP
MB
ACA
CL
CA3
CP
MD
AV
VL no stim
VL sES
5 μA, 8 Hz
5 μA, 28 Hz
5 μA, 140 Hz
rest
run

99

**Figure S7. Spike-phase entrainment at rest and during running.** Vector length (VL) for individual units (gray lines) across brain areas during rest and run locomotion states, shown for baseline (top panel, gray) and sES conditions (bottom panel, colored) at 5 µA for each stimulation frequency (8 Hz, green; 28 Hz, orange; 140 Hz, magenta). Colored lines indicate the median VL per area. Statistical significance of VL differences between rest and run locomotion states shown above each area (●●●● $p \leq 0.0001$; ●●● $p \leq 0.001$; ●● $p \leq 0.01$; ● $p \leq 0.05$; Mann-Whitney U-testing with Benjamini-Hochberg correction for multiple comparisons).

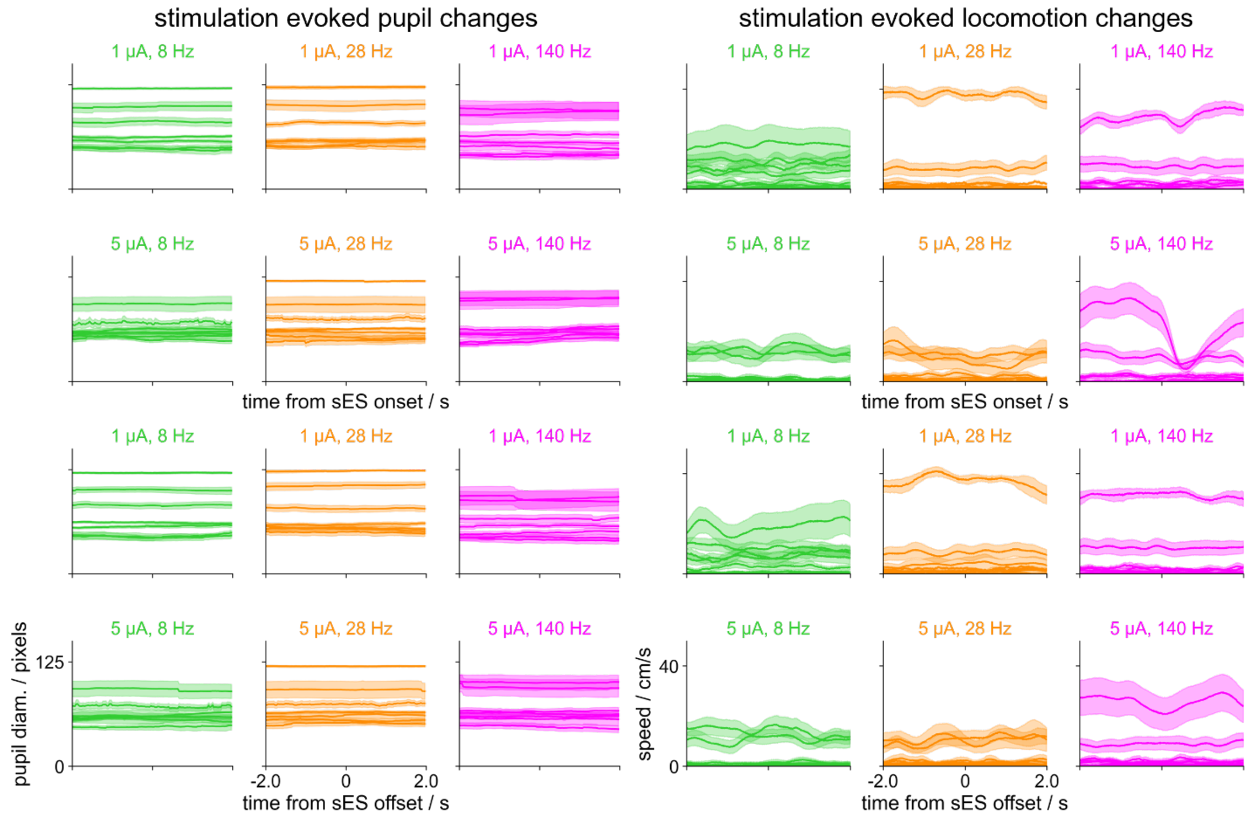


**Figure S8. Stimulation-triggered changes in pupillometry and locomotion.** Stimulation-triggered average of the pupillometry signal (left) and wheel speed (right) around the onset (rows 1–2) and offset (rows 3–4) of each stimulation protocol (1 µA and 5 µA at 8 Hz, green; 28 Hz, orange; 140 Hz, magenta). Each line represents the average signal of one animal (± SEM) in a window of ±2 s around stimulation onset/offset (pupillometry: N=7; wheel speed: N=13). No systematic changes in pupil size or locomotion were observed in relation to stimulation onset or offset.

#### Author contributions

Conceptualization: C.A.A.; Formal analysis: I.R., S-Y.L., A.P., C.A.A.; Investigation: I.R., L.C.M, C.A.A.; Funding acquisition: C.A.A., S-Y.L.; Methodology: I.R., S-Y.L., L.D.C., C.K., C.A.A; Software: I.R., C.A.A.; Writing – original draft: I.R., C.A.A.; Writing – review and edits: all authors.

#### Code Availability

Code is available at the following link: https://github.com/anastassiou-team/NP_brain_stim_INVIVO

#### Data Availability

Data analyzed in this manuscript are publicly available in: https://github.com/anastassiou-team/NP_brain_stim_INVIVO

#### Acknowledgments

We thank the Allen Institute Animal Care, the Lab Animal Services teams for mouse husbandry and care; the Allen Institute Manufacturing and Process Engineering team for experimental hardware and software support; and the Imaging teams. We wish to thank the Allen Institute founder, Paul G. Allen, for his vision, encouragement, and support.

We gratefully acknowledge funding from the National Institute of Neurological Disorders and Stroke of the National Institutes of Health under Award Numbers R01NS120300 and RO1NS130126. The content is solely the responsibility of the authors and does not necessarily represent the official views of the National Institutes of Health.

#### Competing Interests Statement

CK holds an executive position and has a financial interest, in Intrinsic Powers, Inc. C.A.A. is the co-founder of Antikythera Neuro.

#### Online Methods

Experimental procedures closely followed those described in Claar et al. [44] and Russo et al. [26]. A summary of these methods and details of procedures that differ are provided below.

71 Marks, L. C., Rembado, I. & Claar, L. D. . Simultaneous Recording of Neuropixels and EEG in Head-Fixed Mice. *protocols.io* [*https://doi.org/10.17504/protocols.io.14egn9q16l5d/v1*](https://doi.org/10.17504/protocols.io.14egn9q16l5d/v1).
